# Structural variant-based pangenome construction has low sensitivity to variability of haplotype-resolved bovine assemblies

**DOI:** 10.1101/2021.11.02.466900

**Authors:** Alexander S. Leonard, Danang Crysnanto, Zih-Hua Fang, Michael P Heaton, Brian L. Vander Ley, Carolina Herrera, Heinrich Bollwein, Derek M. Bickhart, Kristen L. Kuhn, Timothy PL. Smith, Benjamin D. Rosen, Hubert Pausch

## Abstract

Advantages of pangenomes over linear reference assemblies for genome research have recently been established. However, potential effects of sequence platform and assembly approach, or of combining assemblies created by different approaches, on pangenome construction have not been investigated. We generated haplotype-resolved assemblies from the offspring of three bovine trios representing increasing levels of heterozygosity that each demonstrate a substantial improvement in contiguity, completeness, and accuracy over the current *Bos taurus* reference genome. Diploid coverage as low as 20x for HiFi or 60x for ONT was sufficient to produce two haplotype-resolved assemblies meeting the standards set by the Vertebrate Genome Project. Structural variant-based pangenomes created from the haplotype-resolved assemblies demonstrated significant consensus regardless of sequence platform, assembler algorithm, or coverage. Inspecting pangenome topologies identified 90 thousand structural variants including 931 overlapping with coding sequences; this approach revealed variants affecting *QRICH2, PRDM9, HSPA1A, TAS2R46*, and *GC* that have potential to affect phenotype.

## Introduction

Cattle are a substantial component of global animal-based food production, and are raised for meat, milk, or both. Two subspecies of cattle, taurine and indicine, have emerged from at least two distinct domestication events (Loftus et al., 1994; Pitt et al., 2019)), with artificial selection for production goals or environmental adaptation contributing to diversity within cattle, resulting in the current existence of hundreds of taurine and indicine cattle breeds. Interbreeding and introgressions with other bovids, like yak and banteng (N. Chen et al., 2018; Wu et al., 2018), further drive an increase in genetic diversity within Bovinae.

The *Bos taurus* reference genome was first drafted in 2004 and was based on whole genome shotgun sequence of a Hereford cow, supplemented with sequences of bacterial artificial chromosome clones prepared from DNA of her sire (Elsik et al., 2009). A major revision using Pacific Biosciences continuous long read (CLR) sequencing of the same cow was recently reported (ARS-UCD1.2; (Rosen et al., 2020)) and remains the accepted reference for conducting genomic studies in cattle due to extensive annotation efforts and connections to historical analyses, despite more recent bovine assemblies having higher contiguity and accuracy (Heaton et al., 2021; Oppenheimer et al., 2021; Rice et al., 2020).

Sequence variability between cattle breeds at both the single nucleotide (SNP) and short insertion or deletion (indel) level has been extensively characterized through reference-guided approaches (Daetwyler et al., 2014; Hayes & Daetwyler, 2019; J. Kim et al., 2017). Short sequencing reads have also been used to study larger structural variants (SVs) and variation located in repetitive or challenging regions across Bovinae (Butty et al., 2020; Kommadath et al., 2019; Mei et al., 2020), although their accuracy is limited compared to using long sequencing reads (Mahmoud et al., 2019). SVs may be involved in expression quantitative trait loci more often than previously estimated, and can impact gene expression more than shorter variants (Chiang et al., 2017). However, previous studies suffer from potential reference bias, because the use of a single taurine cattle reference assembly fails to reflect the immense genomic diversity present in global breeds of domestic cattle (Crysnanto et al., 2019, 2021; Crysnanto & Pausch, 2020; Talenti et al., 2021).

Pangenomes have long been proposed (Tettelin et al., 2005) as a way to better reflect variation present in a group of individuals (e.g., breed, species, clade, etc.). Pangenomes can be constructed from variants called through reference-guided approaches (Garrison et al., 2018; Hickey et al., 2020), contigs assembled from reads which failed to align to the reference (Sherman et al., 2018), or multiple whole-genome assemblies (Armstrong et al., 2020; Li et al., 2020). The latter approach may more faithfully capture challenging regions and SVs due to the complexity of calling and representing nested variation. Third-generation sequencing continues to become more cost-effective and accessible, making population-scale *de novo* assemblies more feasible. An influx of high-quality assemblies makes the need for pangenome representations more pressing, although the effects of integrating disparate assemblies into pangenomes are unknown.

The present study combines de-novo assembled genomes from three bovine trios of varying heterozygosity into structural variant-based pangenomes. Haplotype-resolved, reference-quality assemblies are generated from the trio offspring using Pacific Biosciences high-fidelity (HiFi; Eid et al., 2009; Wenger et al., 2019) and Oxford Nanopore Technologies (ONT; Mikheyev & Tin, 2014) sequencing and various assembly algorithms. This set of assemblies from disparate sequencing and origin is then used to evaluate the effects on pangenome construction depending on assembly approach, or when constructed from a combination of approaches. The utility of a bovine pangenome is then demonstrated by analyses using SVs to assess evolutionary relationships between Bovinae, recovering putatively trait-associated SVs, and quantifying SV-coding sequence overlaps.

## Results

The three examined bovine trios (Figure 1a-c) reflect diverse breeding strategies (within-breed, inter-subspecies, and inter-species) and increasing heterozygosity. The first F1 (OxO) was a cross between two Original Braunvieh cattle (Bos taurus taurus), but was still substantially less inbred compared to the cow used for the ARS-UCD1.2 reference (pedigree-based coefficient of inbreeding of 0.07 compared to 0.30 (Rosen et al., 2020) respectively). The second F1 (NxB) was a cross between Nellore (Bos taurus indicus) and Brown Swiss (Bos taurus taurus) cattle and the third F1 (GxP) was an inter-species cross between a gaur (Bos gaurus) bull and a Piedmontese (Bos taurus taurus) cow. HiFi and ONT reads were collected for each F1, and short reads were collected for all animals in the trios (Figure 1d, Supplementary Table 1). F1 long reads were separated into paternal, maternal, and unknown origin (Koren et al., 2018). Separability, the proportion of reads assigned to a parent-of-origin bin, improved significantly from 81.1% to 99.9% with increasing heterozygosity for HiFi reads, but was near perfectly separable for ONT at all examined heterozygosities (Figure 1e, Supplementary Table 2).

**Figure 1.**
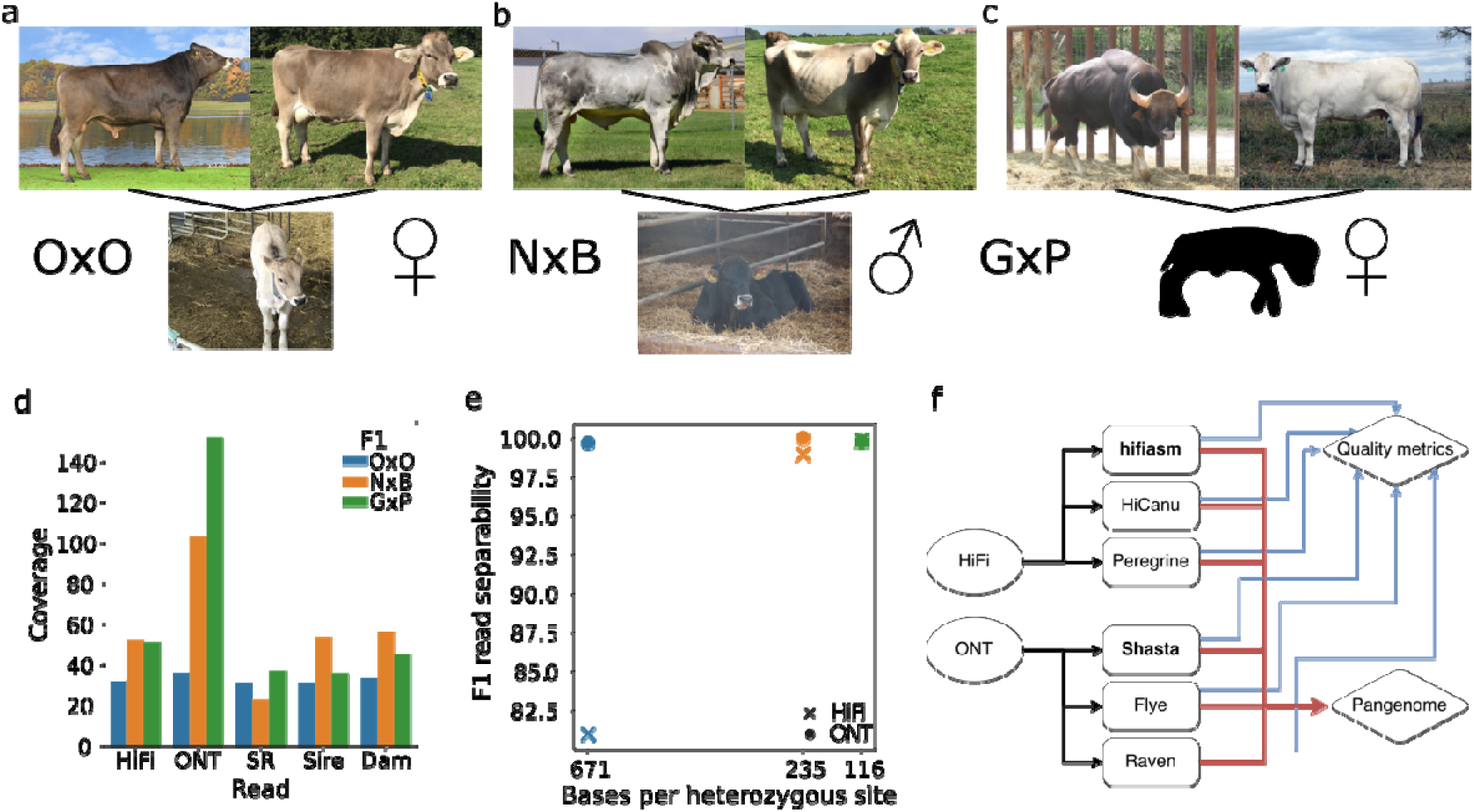
Overview of bovine trios. a) – c) Representative animals for the parents of the three bovine trios and the respective F1s (OxO, NxB, and GxP) examined in this study. The OxO and GxP were female, while the NxB was male. d) The three respective F1s were sequenced to 32-, 52-, and 51-fold HiFi coverage, with read N50 of 20, 21, and 14 Kb. ONT sequencing was performed to 36-, 103-, and 152-fold coverage respectively, with read N50 of 65, 45, and 49 Kb. Coverage is determined with respect to an assumed genome size of 2.7 Gb. F1 short reads (SR) were collected to 31-, 23-, and 37-fold coverage. e) Separating F1 reads into parental haplotype bins improved with increasing heterozygosity for HiFi, but F1 reads were nearly 100% separable for ONT even at low heterozygosity (color from panel d). f) HiFi reads were assembled with hifiasm, HiCanu, and Peregrine, while ONT reads were assembled with Shasta, Flye, and Raven. Bold type indicates the tools used to produce the assemblies that are discussed in detail. Assemblies were assessed individually for quality metrics (blue lines) as well as integrated together into pangenome analyses (red lines).

### Assembling bovine genomes

Investigating the resiliency of pangenome graphs to variable input assemblies was initiated by examining the properties of each type of assembly and their sensitivity to input data parameters. Contigs were assembled for each F1 using multiple assemblers for HiFi and ONT data (Figure 1f, methods). Contigs were scaffolded by alignment to ARS-UCD1.2 to produce the final assemblies, an approach with minimal bias given the well-curated reference and highly contiguous assemblies (Alonge et al., 2019, 2021). The gaur has a centromeric fusion of chromosomes 2 and 28 resulting in a different karyotype than that of domestic cattle (Mamat-Hamidi et al., 2012), but this scaffolding approach was still successful for the paternal haplotype of the GxP (Supplementary Figure 1a). The haplotype-resolved assemblies for the inter-subspecies NxB and inter-species GxP are directly breed/species-specific, while both haplotypes of the OxO constitute the same breed, resulting in six haplotype-resolved assemblies, five of which represent novel breed/species. For breed/species-level analysis, we only use the maternal haplotype of the OxO by choice, while analyses displaying intra-breed variation use both haplotypes (methods). The “quality” of each assembly was assessed by several widely used metrics: contiguity (NG50) representing the size distribution of contigs, phasing (PG50) to characterize haplotype separation, correctness (QV) quantified as Phred-scaled base-error rate, and completeness (BUSCO) which approximates the percentage of near-universal, single-copy genes that were identified. These metrics provide a coarse-grained summary of assembly quality amenable to comparisons.

The assemblies produced by hifiasm for HiFi data and Shasta for ONT data were selected for further analysis based on having the best quality metrics (Table 1) and computational tractability compared to four other tested assemblers (Supplementary Table 3). These assemblies were of reference quality with every examined metric exceeding those of the current *Bos taurus* Hereford-based reference. Improvements over the current reference are substantial, including a 3-11x reduction in autosomal gaps, 1.8-3.6x increase in NG50, and 3-22x reduction in base errors. Furthermore, they all exceed the current standards set by the Vertebrate Genome Project (VGP) (Rhie et al., 2021). The ONT-based assemblies were marginally above the QV targeted by the VGP, but other metrics for these assemblies such as the contig or phased contiguity are orders of magnitude greater than VGP thresholds.

**Table 1.**
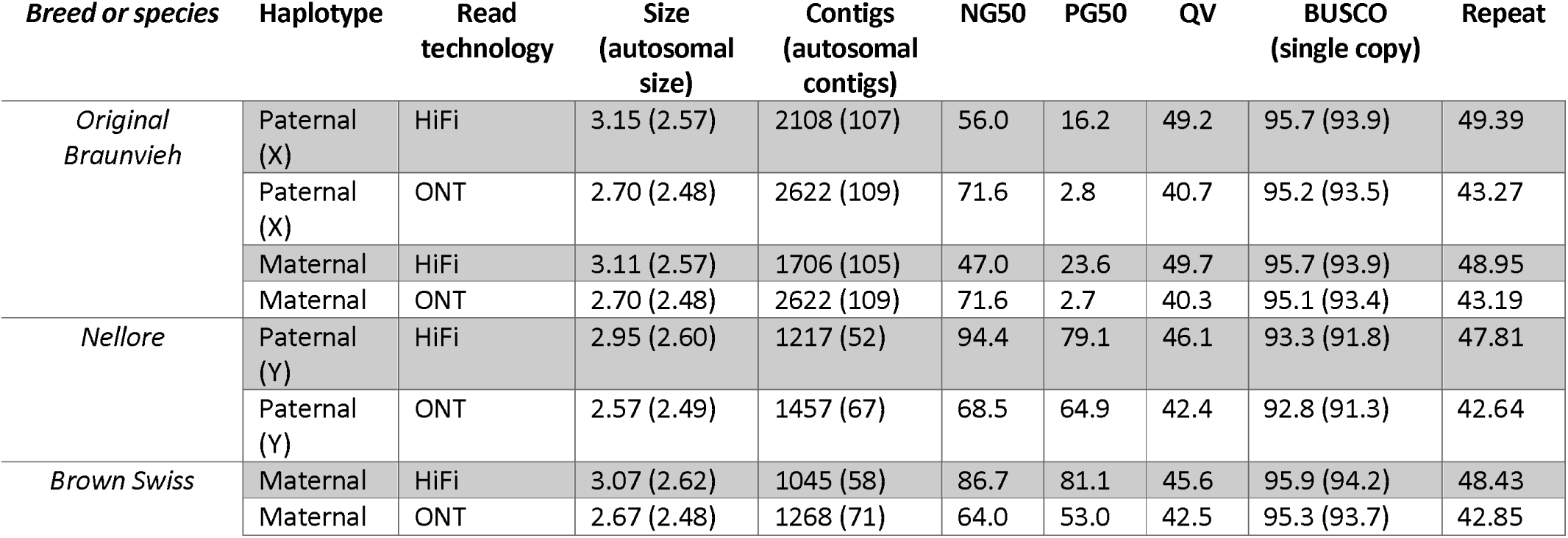
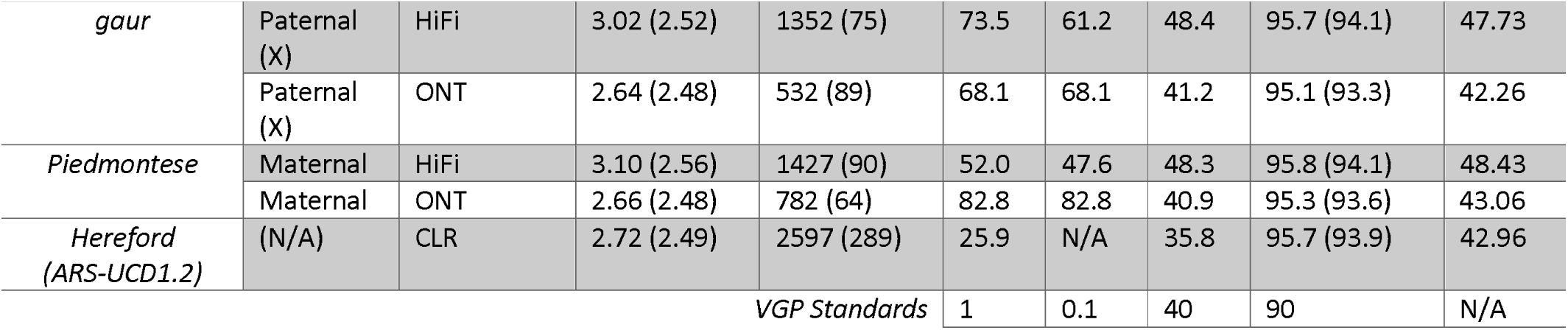
Quality metrics for ten haplotype-resolved assemblies. The assembly haplotype is either maternal or paternal (indicating either an “X” or “Y” paternal sex chromosome). The ARS-UCD1.2 reference is not haplotype-resolved and lacks sufficient parental data to assess phasing, hence the N/A. Size and contigs refer to the entire genome assembly, while the autosomal values only measure chromosomes 1 through 29. NG50 is the contig N50 using the ARS-UCD1.2 reference sequence as the expected length. PG50 is NG50 after splitting contigs into haplotype-phased blocks. Phasing and QV are determined through merqury using parental and F1 short reads. Scaffolded NG50 is not shown, as all assemblies are effectively end-to-end (excluding centromeres and telomeres), with values greater than 100 Mb. Assemblies are available online (Leonard, 2022).

The HiFi- and ONT-based assemblies were generally comparable, however there were notable differences in the average assembly correctness and genome size metrics. Shasta assemblies averaged QV 41.5 after one round each of polishing with ONT reads and short reads, while hifiasm assemblies reach QV 47.6 without any polishing. The log scale of QV means that the hifiasm assemblies had a 4-fold reduction in base errors compared to the Shasta assemblies, indicating the ability of HiFi data to achieve higher quality in fewer steps. In contrast, phasing in Shasta assemblies is better compared to hifiasm, while both platforms showed improved phasing at higher heterozygosity in agreement with the relative ability to sort F1 reads by parental origin prior to assembly. There was insufficient ONT coverage for the OxO to perform the bin-then-assemble approach used for the NxB and GxP and so instead a diploid polishing approach was used to phase the haplotypes (methods), resulting in reduced PG50.

The mean autosomal genome size of the assemblies generated by hifiasm and Shasta was 2.57 ± 0.03 Gb and 2.48 ± 0.004 Gb respectively, such that on average each hifiasm autosome was 2.8 Mb longer than ARS-UCD1.2, and 0.34 Mb shorter for Shasta. The additional length of hifiasm autosomes was primarily due to the presence of more repetitive sequences (43.5% repetitive content in hifiasm versus 41.7% in Shasta), especially centromeric repeat sequence which accounted for 56% of the additional sequence compared to Shasta assemblies. Previous studies have shown that the higher accuracy of HiFi reads allows assemblers to confidently assemble through more repeats despite having shorter read lengths (Chu et al., 2021), leading to extension of autosomal contigs into flanking centromere sequence. Hifiasm assemblies also contained more sequence in contigs not assigned to chromosomes (average of 300 Mb) compared to Shasta assemblies (50 Mb). These unassigned contigs were composed primarily of repetitive sequences (Supplementary Table 4) including novel centromeric sequence and long terminal repeats. Unplaced contigs were generally higher in repeat content than the scaffolds (88% versus 48% for hifiasm and 87% versus 42% for Shasta), and thus would present a challenge to scaffolding by any technology including the reference alignment approach applied here.

Bovine autosomes are acrocentric (Blazak & Eldridge, 1977), and so a complete bovine assembly is conceptually closer to “centromere-to-telomere”. Hifiasm assemblies contained substantially more centromeric sequence than Shasta assemblies, respectively averaging 2.01 and 0.14 Mb per autosome, compared to 0.08 in the ARS-UCD1.2 reference (Figure 2a). Similarly, hifiasm autosomes average 2.6 Kb of vertebrate telomeric repeats (TTAGGG) within 10 Kb of the chromosome end, compared to 0.8 Kb for the ARS-UCD1.2 reference. Telomeres were almost entirely missing in Shasta assemblies, averaging only 88 bp of telomeric repeats per autosome (Figure 2b), a reported issue with ONT basecalling (Tan et al., 2022). Chromosomes which contain at least 50 kb of centromeric repeats at the proximal end and 500 bp of telomeric repeats at the distal end are considered to be end-to-end (but not necessarily “complete”), of which there were 5 for ARS-UCD1.2, and a mean of 13.2 and 1.2 for hifiasm and Shasta across the five breeds/species. Hifiasm and Shasta assemblies had a near equal distribution of gaps, averaging about 1.5 gaps per autosome, compared to nearly 9 in the ARS-UCD1.2 reference (Figure 2c). These observations hold in general for all HiFi- and ONT-based bovine assemblies investigated (Supplementary Figure 2). These differences are visible on the example Brown Swiss chromosome ideograms in Figure 2d. Both HiFi- and ONT-based assemblies were able to assemble the 16 Kb bovine mitochondrial DNA (Anderson et al., 1982).

**Figure 2.**
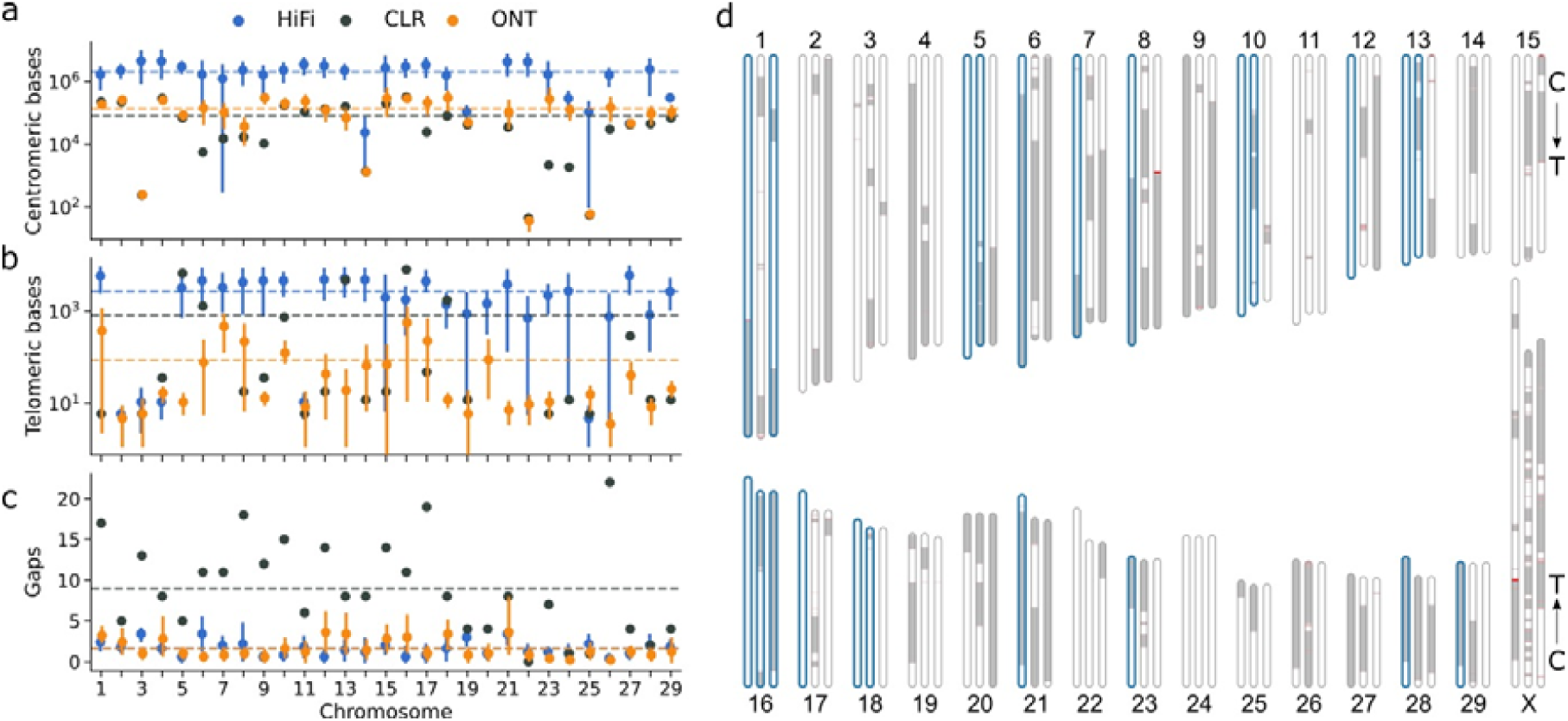
Centromeric and telomeric completeness of assemblies produced by hifiasm and Shasta. a) The mean number of bases identified per autosome as “Satellite” by RepeatMasker for the five hifiasm (blue) and five Shasta (orange) assemblies, where error bars indicate the 95% confidence interval. The black dots represent values from the CLR-based ARS-UCD1.2. Dashed lines indicate the autosome-wide mean for the respective color of points. Mean values of 0 (e.g., chromosome 20) are not shown due to the log scale. b) Similar to a), but the number of bases in telomeric repeats within 10 Kb of chromosome ends. c) Similar to a), but the number of scaffold gaps. d) Chromosome ideograms for ARS-UCD1.2 (center), and Brown Swiss assemblies produced by hifiasm (left) and Shasta (right). Scaffolded contigs alternate white/grey across gapped regions, which are colored red. Chromosomes which are predicted to extend from centromere to telomere are bolded in blue. Arrows indicate the centromere-to-telomere directionality of the chromosomes.

### Optimal sequencing coverage depths

Impact of sequencing depth on assembly quality was assessed using the Brown Swiss haplotype of the NxB as an example. Subsets of the 52x diploid HiFi and 55x haploid (trio binned) ONT reads, respectively, were randomly sampled to mimic lower sequencing depths (Figure 3). Completeness metrics (e.g., BUSCO or k-mer content) plateau when coverage increased above 25-fold, however other metrics like contiguity or correctness continued to benefit from higher sequencing coverage with diminishing returns. The trio aware mode of hifiasm only required about 19x diploid coverage of HiFi reads to meet the VGP targets. Shasta required around 28x haploid coverage (corresponding to roughly 56x diploid coverage) to achieve the necessary QV, although 17x haploid (34x diploid) coverage fulfills the contiguity and completeness targets. The minimum VGP-satisfying coverage varies slightly for the different F1s due to different input sequencing read properties but is approximately consistent across all examined bovine trios (Supplementary Figure 3).

**Figure 3.**
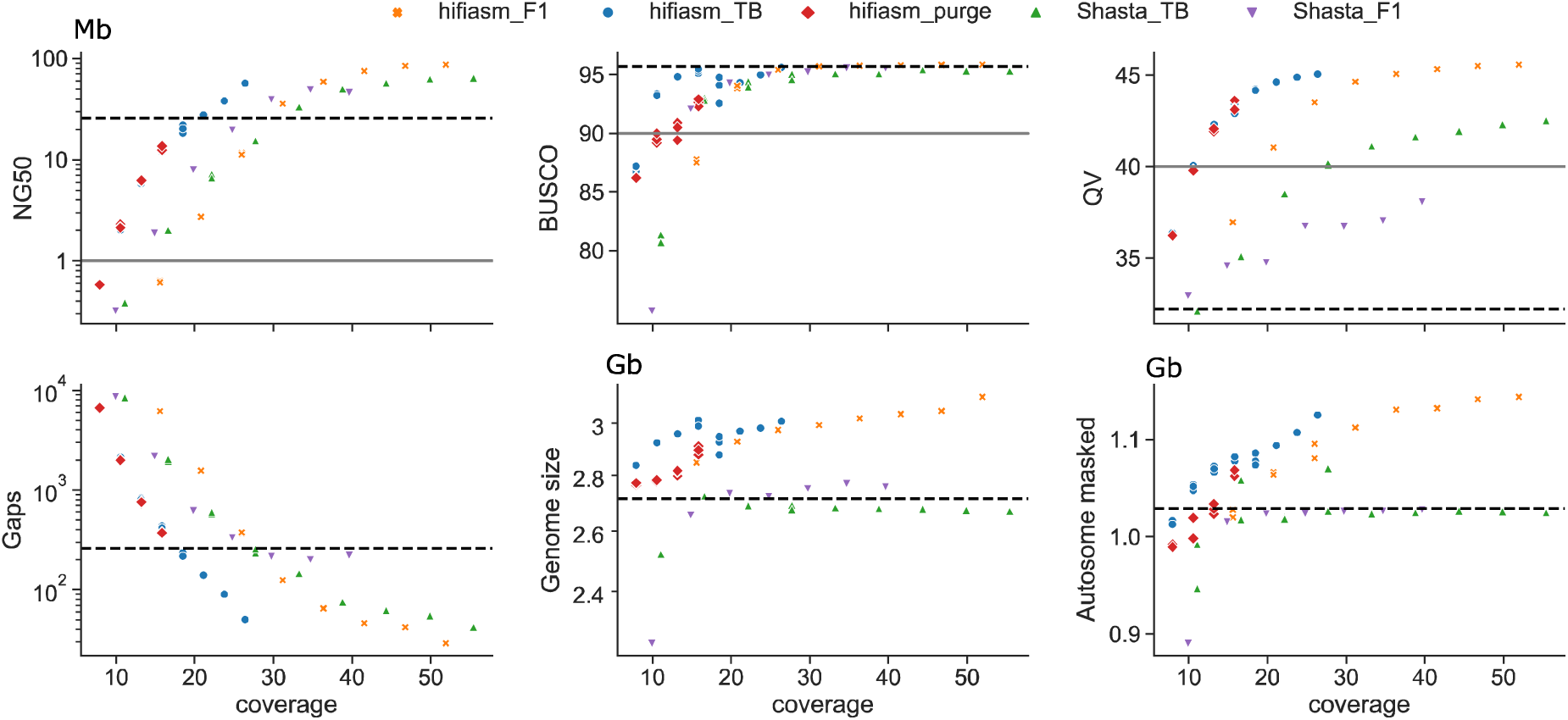
Assembly quality at subsampled coverages. Trio aware hifiasm (hifiasm_F1) uses diploid coverage while Shasta (Shasta_TB) uses haploid coverage. We additionally examined trio binned hifiasm (hifiasm_TB) using haploid coverage and the polish-phased Shasta approach using diploid coverage (Shasta_F1). NG50, BUSCO, QV, autosomal gaps, and genome size are defined in Table 1, while autosomal masked is the number of autosomal bases within repetitive elements as identified by RepeatMasker. The black dashed line represents the relevant value for the ARS-UCD1.2 reference and the grey solid line is the VGP target where applicable. Three subsampling replicates were performed for lower coverage assemblies (<20x for HiFi and <30x for ONT) due to their higher stochasticity. For trio binned hifiasm assemblies below 15x coverage, we manually set the duplication purging parameter (hifiasm_purge) and reran on the same subsamplings.

The higher accuracy of HiFi reads allows hifiasm to exploit sequence common to both haplotypes during trio aware assembly. Reaching a comparable quality through a trio binned approach required approximately 16% higher coverage, with 11x haploid coverage (22x diploid) necessary. The higher error rate of ONT reads makes haplotype-aware correction and phasing challenging, and so diploid assembly followed by haplotype separation or diploid-aware polishing is less effective than trio binning (Figure 3, up triangles versus down triangles). Phasing is particularly poor, with the PG50 25 times smaller on average compared to trio binning approaches.

There is an increased risk of coverage gaps when sequencing coverage is reduced, even if the resulting assemblies achieve certain genome-wide standards. When 11x HiFi coverage is aligned to ARS-UCD1.2 and binned into 10 Kb windows, there are 950 regions on average of near total dropout (<1x coverage) across the autosomes. This drops by 23% at 13.5x coverage and by 30% at 16x coverage, as the effects of stochasticity are reduced. While the overall assembly quality does not fluctuate substantially between random subsamplings (Figure 3), it may overestimate the quality at specific regions. Furthermore, default parameters of assemblers are typically tuned to higher coverage and assembling at low coverage can introduce subtle issues. Hifiasm could underestimate a parameter related to duplication purging at coverages below 15x, resulting in a sharp transition to larger assemblies with more duplicated BUSCO genes (Figure 3, blue circles versus red diamonds). Manually setting the parameter to its expected value recovered similar behavior seen in higher coverage assemblies.

### Constructing a bovine pangenome

Sequencing technologies and assemblers evolve rapidly, and so even recently generated bovine assemblies, including the ones reported here, have been produced under non-uniform conditions (e.g., (Crysnanto et al., 2021; Talenti et al., 2021)). Given the differences we observed between HiFi- and ONT-based assemblies, especially in comparison to the CLR-based ARS-UCD1.2 reference, it was crucial to examine how pangenome construction responded to different assembly inputs.

Pangenomes were constructed with minigraph (Li et al., 2020) using ARS-UCD1.2 as the initial backbone of the graph structure. Assemblies were iteratively added into the graph, and regions of synteny were ignored while sufficiently diverged subsequences (>50 bp) instead augmented the graph with new nodes (“bubbles”). Pangenomes for each autosome were constructed from all hifiasm assemblies, all Shasta assemblies, or random mixtures of the two. Graph properties of each pangenome in terms of the amount of non-reference sequence added, were generally robust to the input assemblies (Figure 4a,b, Supplementary Table 5). Pangenomes constructed from five hifiasm assemblies had more non-reference sequence added compared to five Shasta assemblies (82.5 Mb across 88.5k nodes versus 63.5 Mb across 90.2k nodes), in agreement with the greater completeness observed in hifiasm assemblies. Approximately 92.1% of identified SVs were common between hifiasm and Shasta pangenomes, with 3.6% and 4.3% unique to each respectively (Figure 4c). There was not a clear bias between HiFi- and ONT-based pangenomes, with only 1.8 and 0.39 Mb non-reference sequence uniquely identified in each. These bubbles were more repetitive than autosome-wide averages (53% and 45% respectively, Supplementary Table 6). Minigraph is sensitive to the order of integration, and on rare occasions constructed significantly different bubbles, particularly for palindromic sequences, resulting in larger variance on chromosomes 7 and 12 (Supplementary Figure 4). Pangenomes constructed from lower-coverage assemblies remained robust. We selected hifiasm and Shasta assemblies generated with an average of 21.6x diploid and 24.9x haploid coverage respectively, which approximately satisfy the VGP standards. The hifiasm assemblies have a higher average QV compared to the Shasta assemblies (41.9 versus 39.3), but lower NG50 (2.3 Mb versus 19.5 Mb) and more autosomal gaps (1869 versus 323) (Supplementary Table 7). Although these assemblies are substantially worse than their full-coverage counterparts, the resulting pangenomes are similar (Figure 4a-c). Taking the high coverage SVs as the truth set, the low coverage hifiasm and Shasta pangenomes have an F1 score of 98.5 and 94.7 for SV discovery respectively. The low coverage Shasta pangenomes tend to identify a greater number of SVs not present in other pangenomes, and so may be false positives.

**Figure 4.**
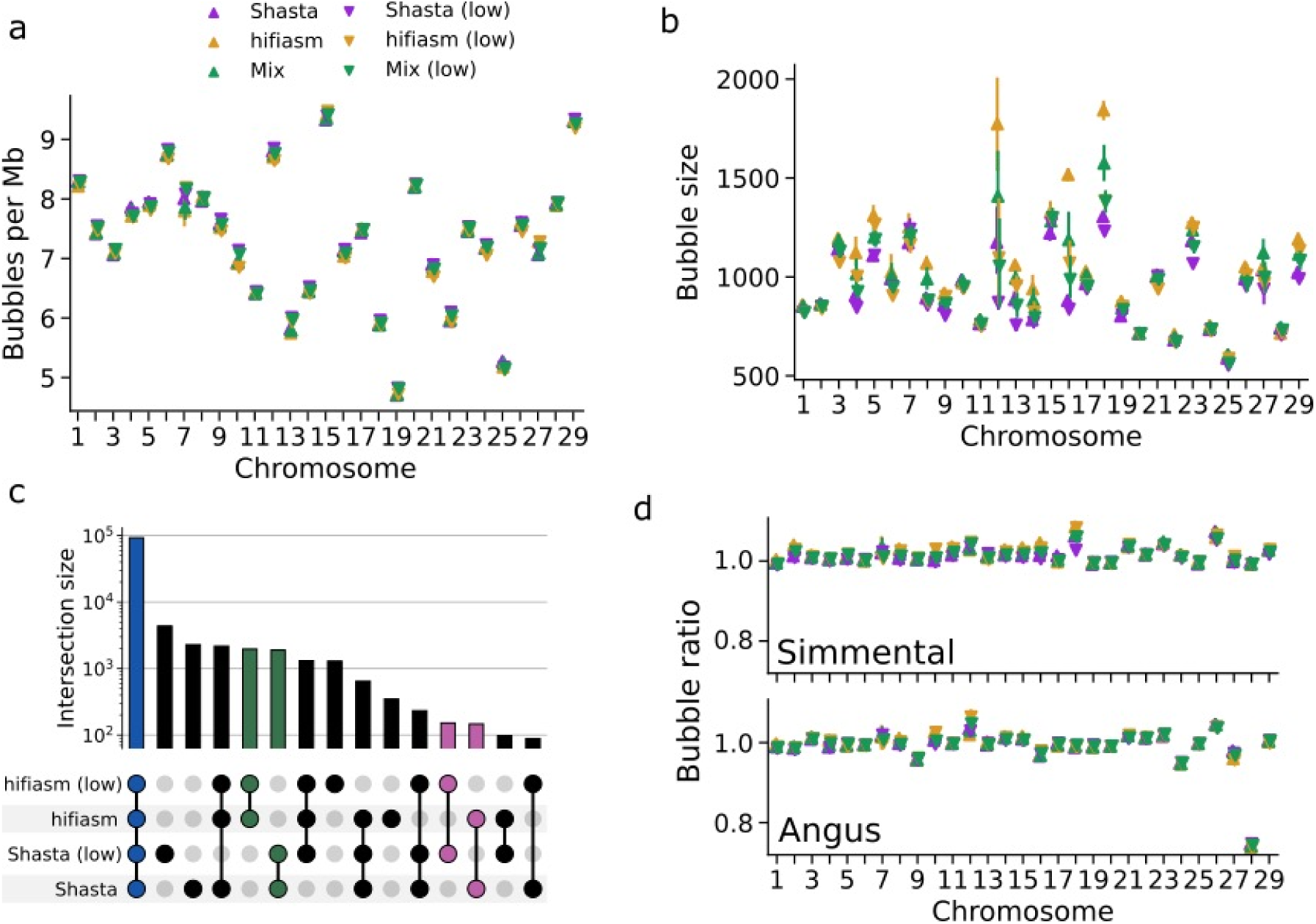
Pangenomes are generally robust to different input assemblies. a) The number of large (>1 Kb) bubbles is highly consistent across hifiasm, Shasta, and mixed pangenomes at both full (up triangles) and lower (down triangles) coverages. b) The mean bubble size is also consistent across different inputs, but bubbles are larger on average in hifiasm pangenomes compared to Shasta pangenomes. c) The vast majority (84.5%) of SVs identified through minigraph are present in all pangenomes (blue). SVs unique to either hifiasm or Shasta (green) only account for about 3.5% of all SVs, while SVs only identified through either full or lower coverage pangenomes are negligible (pink). d) Comparing the number of bubbles present in Simmental-or Angus-backed pangenomes to the ARS-UCD1.2-backed pangenome in a) shows consistency. Angus chromosome 28 is the only exception due to its incomplete sequence. All points reflect the mean over 20 stochastic pangenome constructions, and error bars indicate the 95% confidence interval.

Pangenomes constructed using the existing Angus (CLR-based) (Low et al., 2020) or Simmental (ONT-based) (Heaton et al., 2021) assemblies as backbones (Figure 4d) produced similar results compared to using the Hereford-based ARS-UCD1.2. More non-reference sequence was identified in the lower-quality Angus-backed pangenome (+13%), while the more complete Simmental-backed pangenomes had less (−6%). Reference-bias propagates through minigraph’s pangenomes, such as the missing sequence in Angus chromosome 28 (Lloret-Villas et al., 2021) resulting in 25% fewer bubbles compared to using ARS-UCD1.2 (Figure 4d).

### Quantifying bovine structural diversity through the pangenome

Potential effects of particular assembly approaches or sequencing technologies on the utility of pangenome graphs for structural variation detection were investigated through the analysis of mutual and private bubbles. Structural variation bubbles were associated with their source assembly by retracing each “haplotype walk” through the graph. The phylogenetic topology of their evolutionary relationship was then estimated by counting the number of mutually exclusive bubbles any two assemblies have to construct a condensed distance matrix. The results are consistent with expectations, with the gaur (*Bos gaurus*) largely separate, followed by the Nellore (*Bos taurus indicus*), and then the three taurine cattle (Figure 5a). All constructed pangenomes unequivocally predicted the first two branches, while there was also good agreement within the closely related taurine cattle (Figure 5b).

**Figure 5.**
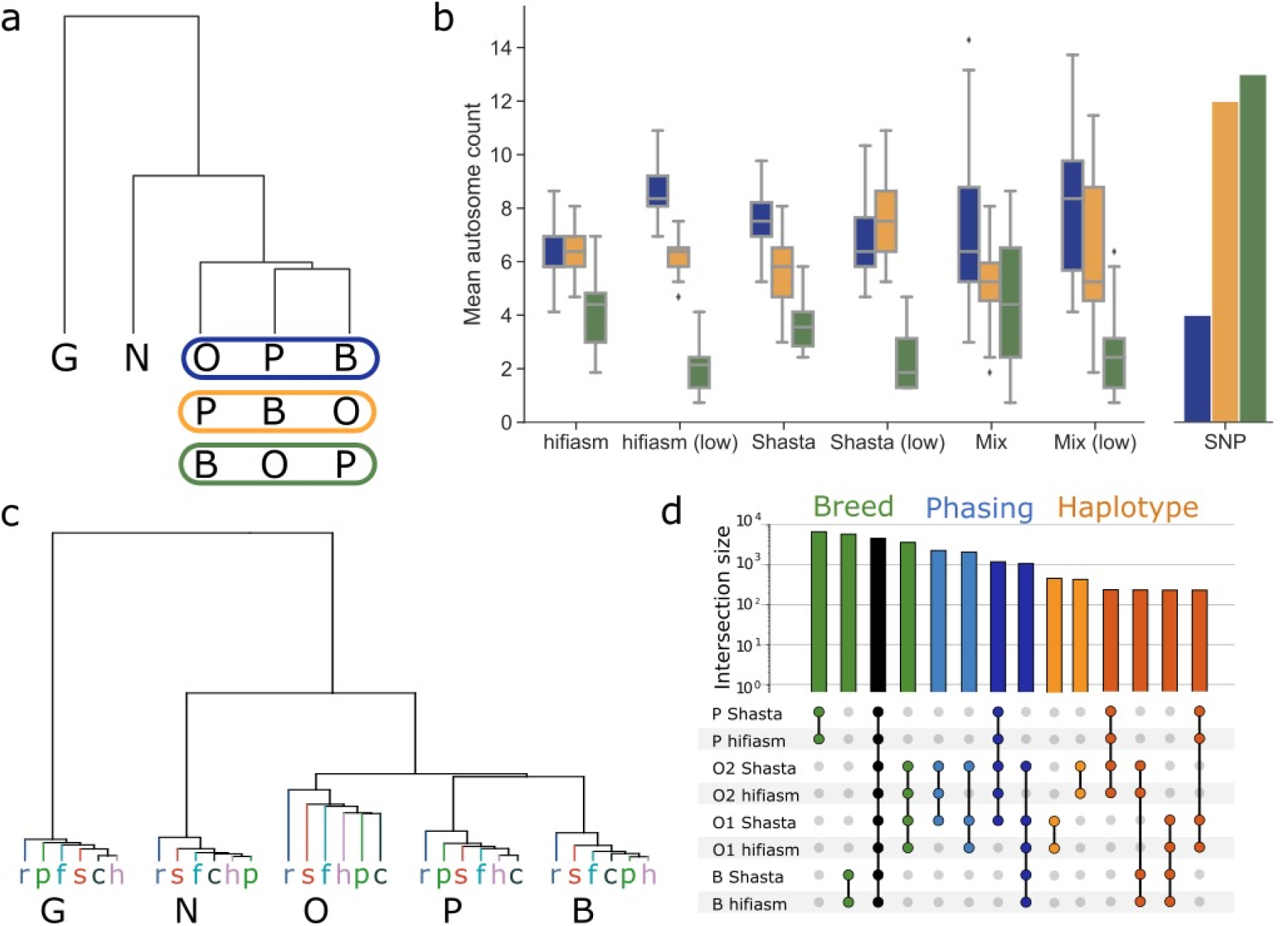
SV-based dendrograms. a) All dendrograms followed the same overall topology, with gaur (G) and Nellore (N) clearly differentiated while the taurine cattle displayed three possible arrangements, with either the Original Braunvieh (O), Piedmontese (P), or Brown Swiss (B) more distantly related. The colored boxes represent the three possible orderings of the taurine cattle. b) Box plots of 20 randomly constructed pangenomes with either all hifiasm, all Shasta, or mixing hifiasm and Shasta assemblies, as well as the low coverage equivalents show good agreement on autosomes displaying a specific topology (color from panel a). The box plots represent the median (center line), first and third quartile (box bottom and top), and 1.5x the interquartile range (whiskers). Outliers beyond this range are marked by diamond markers. The SNP dendrogram, based on parental short reads, generally predicted different topologies. c) Pangenomes including all 30 assemblies from hifiasm (h), Shasta (s), Peregrine (p), Flye (f), HiCanu (c), and Raven (r) predict the same overall topology without ONT or HiFi specific branches. d) An UpSet plot of a taurine cattle pangenome (Piedmontese, Brown Swiss, and the paternal [O1] and maternal [O2] haplotypes of Original Braunvieh) reveals inter-breed variation (green) as well as intra-breed variation in the Original Braunvieh haplotypes (red & orange). We can also identify phasing error candidates in the Original Braunvieh Shasta assemblies, where SVs are common to both Shasta assemblies but not both hifiasm assemblies (light and dark blue).

Several chromosomes (e.g., 2, 6, 9, etc.) are observed to only predict a single taurine topology, even across different assembly inputs and coverages, while other chromosomes (e.g., 1, 12, 16, etc.) have multiple predicted topologies with similar frequencies. This may indicate that certain chromosomes harbor greater structural variation between specific taurine breeds, which may be reflected in phenotypic differences. We compared against a conventional approach, calling small variants from the parental short reads corresponding to the five haplotypes against ARS-UCD1.2 (Figure 5b). While the overall topology and magnitude is similar to that found through the pangenome, there was no significant concordance within the taurine cattle, as might be expected given the low linkage disequilibrium between small and structural variants (Yan et al., 2021).

Genome level dendrograms were mostly consistent across different combinations of inputs, demonstrating that pangenome robustness extends beyond graph properties and into applications. The same general topology was recovered for all examined assemblers, both individually and combined, as well as the lower coverage assemblies. The dendrogram correctly places all assemblies of each breed on their own branch (Figure 5c) when 31 pangenome assemblies are included (1 reference backbone plus 5 breeds/species x 6 assemblers). Some HiFi- and all ONT-based assemblies for Original Braunvieh are only partially phased due to limited coverage, resulting in greater structural variation within these assemblies (Figure 5d). The variation between the parental haplotypes of the OxO results in the higher branching point for “O” in Figure 5c, and highlights the importance of cleanly resolved haplotypes for downstream analyses.

### Pangenome topology at trait-associated SVs

There presently is no “truth set” of bovine structural variants appearing in the haplotypes represented to benchmark the success of each pangenome graph. Therefore, we selected several previously identified SVs known to impact phenotype which were present in our pangenomes for further investigation. One case is a multi-allelic copy number variation (CNV) at 86.96 Mb on BTA6 encompassing an enhancer of the group-specific component (*GC*) gene. This CNV has pleiotropic effects on mastitis resistance and other dairy traits, and segregates in different breeds of cattle (Lee et al., 2021; Olsen et al., 2016), although the prevalence of the CNV has not yet been determined for the breeds/species included in our bovine pangenome. All examined assemblers predicted duplication of a 12 Kb segment in Brown Swiss and Original Braunvieh (Figure 6a,b, Supplementary Figure 5), although hifiasm required reassembling a small subset of reads as the duplication was missing in the original assembly (methods). The *de novo* assembled duplicated segments contained various repetitive elements (Figure 6c) matching those previously reported (Lee et al., 2021). There was poor consensus in copy number among the assemblies (Supplementary Figure 5). Inspection of coverage data retrieved from HiFi- and ONT-binned long read alignments against the ARS-UCD1.2 reference confirmed that the gaur, Piedmontese, and Nellore haplotypes have no additional copies, while the Brown Swiss and Original Braunvieh haplotypes harbor between 2 and 4 additional copies of the 12 Kb segment (Figure 6d, Supplementary Notes). Coverages derived from the NxB and OxO F1 short read alignments are consistent with these values but are unable to resolve the haplotype-specific copy number disagreement observed between HiFi and ONT coverage, as well as between alignments and assemblies (Figure 6e).

**Figure 6.**
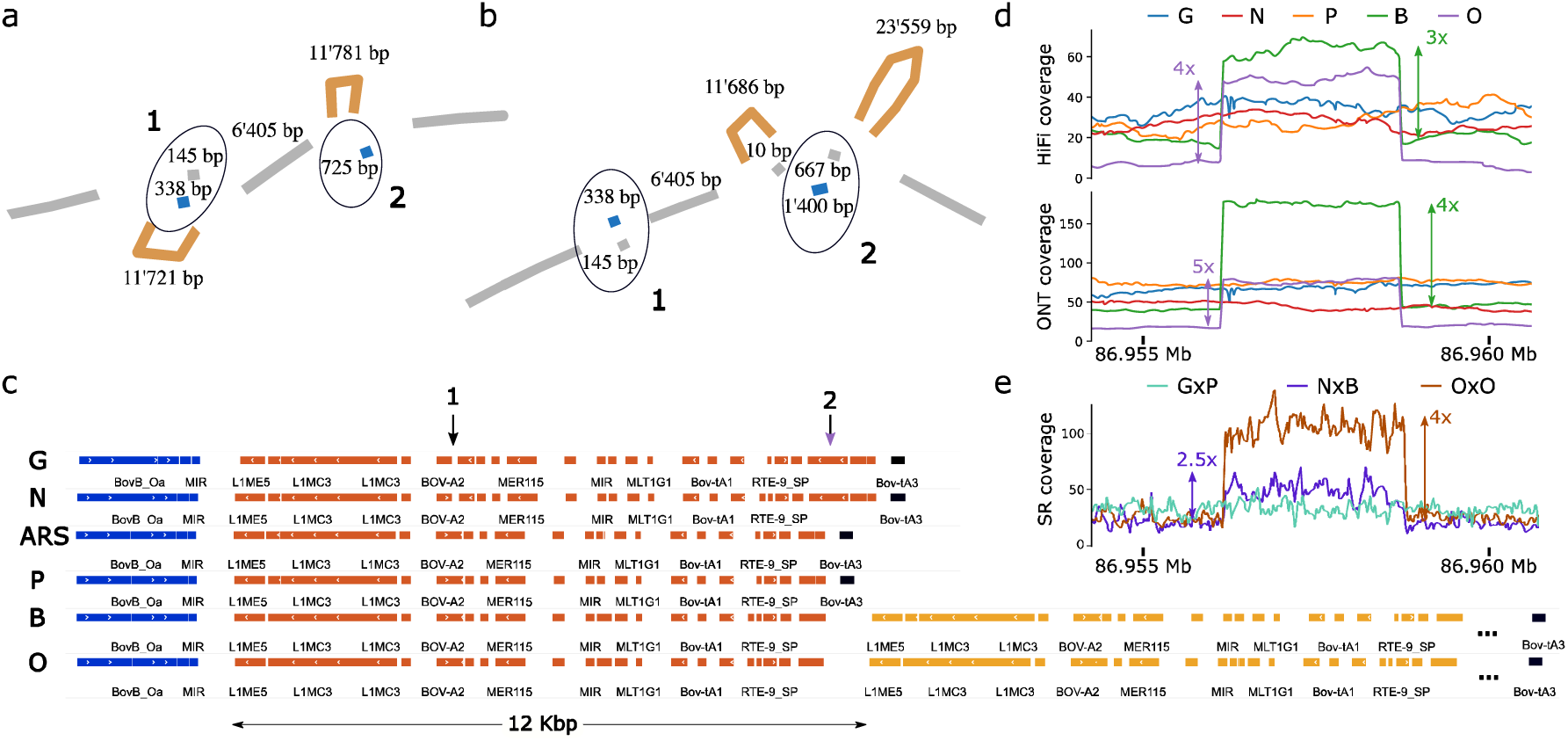
Topology of a tandem duplication on BTA6. a-b) Example subgraphs of the promoter region of GC from a) hifiasm- and b) Shasta-based pangenomes respectively. Reference paths (including those in bubbles) are colored gray, while the tandem duplications are orange. Two insertions observed uniquely in Nellore and gaur haplotypes are blue, shown in circles 1 and 2. Complex bubbles generally have suboptimal topologies due to the lack of base-level alignment. For example, the 725 bp insertion is obvious in a), but appears as the difference between a 1,400 bp and 667 bp path in b). However, both subgraphs identify the approximately 200 bp (1) and 700 bp (2) insertions in Nellore and gaur, as well as the tandem duplication in Brown Swiss and Original Braunvieh. c) The 12 Kb repeat structure (orange) is clearly identified by RepeatMasker across all assemblies, shown here for the ARS-UCD1.2 reference and Shasta assemblies for gaur, Nellore, Piedmontese, Brown Swiss, and Original Braunvieh. The two marked gaur/Nellore insertions (1&2) are consistent with the pangenomes in a-b). One additional copy in Brown Swiss and Original Braunvieh is shown (yellow), while the tandem duplication eventually ends with a similar repeat (Bov-tA3, black) to the other assemblies. d) The identified CNV region shows clear coverage increase in only Brown Swiss and Original Braunvieh, across both HiFi and ONT haplotype-resolved reads, although the HiFi reads suggest one less additional copy than ONT reads. e) F1 short reads also show increased coverage for the NxB and OxO trios. The NxB coverage increase is consistent with only the Brown Swiss haplotype carrying additional copies.

An SV at the *ASIP* gene, encoding the Agouti-signaling protein involved in mammalian pigmentation and located on chromosome 13, was also investigated. This genomic sequence harbors alleles associated with coat color variation in many species including cattle (Girardot et al., 2006; Trigo et al., 2021). Both Nellore and Brown Swiss cattle present variability in coat color ranging from near white to almost black. The NxB was born with a light coat, which darkened as the bull aged (Supplementary Notes). Our pangenome confirmed great allelic diversity between the bovine assemblies involving insertions and deletions of repetitive elements upstream of *ASIP*. However, the previously described variants associated with coat color variation (Trigo et al., 2021) were not in the pangenomes. Short read alignments of the Nellore sire confirmed it carried the previously described SV in the heterozygous state, but the F1 inherited the other haplotype (Supplementary Notes). Thus, the darkening of the coat we observed in the NxB is not due to previously described *ASIP* alleles.

We identified 808 and 845 genes in the ARS-UCD1.2 genome annotation (Refseq release 106) whose coding regions overlap bubbles with complete haplotype information in hifiasm and Shasta based pangenomes respectively, with 722 common to both (Supplementary Figure 6, Supplementary Table 8). This overlap is less than expected under random distribution of the same bubbles (one-sided Wilcoxon test: p=6.9e-10, methods), as expected given the greater evolutionary conservation of genic regions. Out of 229 affected genes which had pLI scores in human orthologues (Fuller et al., 2019), 35 and 36 overlapping bubbles in hifiasm and Shasta pangenomes (30 common to both) exceeded the threshold of 0.9 indicating they are intolerant to change (Supplementary Table 8). Separately, five and three genes that overlapped with bubbles in hifiasm and Shasta pangenomes are listed in the OMIA (Online Mendelian Inheritance in Animals) database of 168 genes that cause phenotypic variation in cattle (Supplementary Table 8); for instance, both hifiasm and Shasta pangenomes contained a bubble encompassing a 36 bp deletion in *ACAN* resulting in the in-frame deletion of 12 amino acids in the gaur assemblies. Variants in *ACAN* are associated with variation in stature in cattle (Cavanagh et al., 2007) and other species (Gibson & Briggs, 2016). The pangenome also recovered the insertion of an 11 *Kb* segment on chromosome 23 in all assemblies. This segment encompasses *HSPA1B* (Hess et al., 2018) which is not annotated in ARS-UCD1.2 (Suqueli García et al., 2017). This 11 Kb insertion is challenging to identify by inspection of short and long read alignments, mainly appearing as elevated coverage over the 2294 bp segment encompassing *HSPA1A*, indicating a similarly sized duplication, with soft-clipped bases extending on both sides (Supplementary Note).

We also investigated genes in this set previously reported to harbor variants that affect phenotype in cattle. For instance, the coding sequence of *QRICH2* overlapped with multiple bubbles indicating tandem duplications of a 30 bp region of the fifth exon (Figure 7a). Loss of function alleles in mammalian *QRICH2* orthologs lead to multiple morphological abnormalities of the sperm flagella (Hiltpold et al., 2021; Shen et al., 2019). We find that the fifth exon of *QRICH2*, which is affected by the coding sequence expansion, is transcribed in high abundance (>30 transcripts per million) in testes of mature taurine bulls (Supplementary Notes). Inspection of long read alignments confirms an expansion of the coding sequence relative to ARS-UCD1.2 that extends the high molecular weight glutenin subunit of the protein by 10 amino acids in our taurine cattle, 60 amino acids in Nellore, and 50 amino acids in gaur (Figure 7b,c). This SV is challenging to resolve with short reads (Supplementary Notes). Another example of potential trait-associated SV lies in *TAS2R46*, related to bitter taste receptors and associated with adaptation to dietary habitats (Dong et al., 2009), which overlapped with a 17 Kb deletion in gaur (Figure 7d,e). This deletion also spanned ENSBTAG00000001761. A final example is found in *PRDM9*, the only known speciation gene in mammals and known to harbor alleles associated with variation in meiotic recombination within and between Bovinae (Ahlawat et al., 2016; Sandor et al., 2012; Zhou et al., 2018), where an SV overlapped with copy number variation in the zinc finger array domain (Figure 7f,g). The Nellore and gaur assemblies contained one zinc finger less, while the paternal haplotype of OxO carried one more relative to ARS-UCD1.2. The maternal haplotype of OxO contained the same number of zinc fingers as ARS-UCD1.2, supporting the intra- and inter-breed/species variation observed for *PRDM9*.

**Figure 7.**
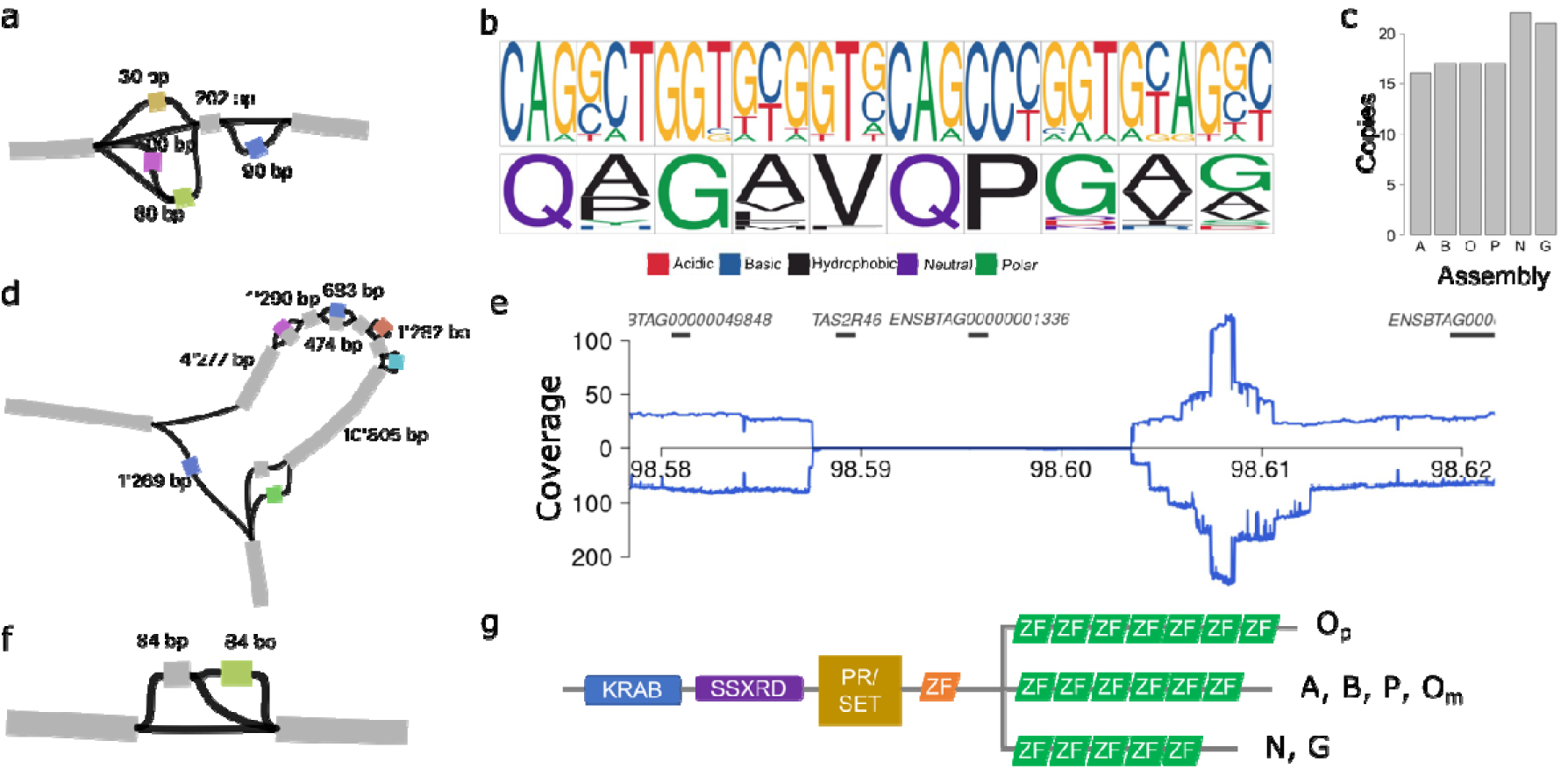
Identification of structural variation in coding sequences of QRICH2, TAS2R46 and PRDM9 through the HiFi-based pangenome. a) Pangenome topology in the fifth exon of bovine QRICH2 revealed tandem repeats of 30 bp sequence. b) Nucleotide (upper) and protein (lower) sequence logo plot of the repeat motif. c) While the ARS-UCD1.2 reference sequence contains 15 copies of the repeat motif, the pangenome revealed 1, 5, and 6 additional copies in the five haplotype-resolved assemblies (A – ARS-UCD1.2, B – Brown Swiss, O – Original Braunvieh, P – Piedmontese, N – Nellore, G – gaur). d) Representation of a 17 kb deletion on BTA5 encompassing TAS2R46 and ENSBTAG00000001761. e) Coverage of binned HiFi (above horizontal line) and ONT (below horizontal line) long read alignments in gaur indicate a large deletion between 98,587,384 and 98,604,401 bp. f) Pangenome topology at the eleventh exon of PRDM9 indicating paths with gain and loss of 84 bp sequence. g) Representation of the domains of PRDM9 in the haplotype-resolved assemblies including a variable number of zinc fingers (ZF) in the different assemblies, where O_m_ and O_p_ are the maternal and paternal haplotypes of the OxO.

## Discussion

Haplotype-resolved assemblies for cattle and related species were constructed using recent sequencing and assembling technologies. Assemblies produced by hifiasm (HiFi) and Shasta (ONT) were substantially more contiguous and correct than those produced by other tools evaluated (Supplementary Table 3) and the current Bos taurus reference sequence, which is a haplotype-merged assembly based on older CLR technology. The higher accuracy of HiFi reads was found to be generally more advantageous to quality measures of assembly and increased completeness of centromeric and telomeric regions compared to the longer but higher-error ONT reads. HiFi-based assemblers also required less compute and storage resources compared to ONT-based assemblers; producing haplotype-resolved hifiasm assemblies required approximately 600 CPU hours and 200 GB of peak memory usage, while the equivalent Shasta (plus polishing) assemblies took 2200 CPU hours and 750 GB of peak memory usage. Correct-then-assemble approaches like Canu (Koren et al., 2017) can be practical for smaller genomes (Wick & Holt, 2021), but on gigabase-sized mammalian genomes like in Bovinae we observed >20 Tb of peak temporary storage and >25k CPU hours for correcting only 30-fold ONT reads. Even recent reference-guided correction approaches like Ratatosk (Holley et al., 2021) still needed approximately 15k CPU hours to correct 55-fold ONT reads. Cutting-edge sequencing and bioinformatic improvements (Baid et al., 2021; Silvestre-Ryan & Holmes, 2021), like the ONT Guppy5 basecaller, will likely assist more efficient assembly, resulting in higher QV and reduced computational load; however, currently the ONT specific requirements might be computationally prohibitive, especially when assembling many samples. Currently there are no approaches to co-assembling HiFi and ONT data that fully exploit the different advantages of the reads; however, HiFi and ONT assemblies can be merged *post hoc* (Alonge et al., 2021) which may improve contiguity (van Rengs et al., 2021), although we observed only minor increases (Supplementary Table 8).

The phased assembly graph approach of hifiasm is most efficient with lower heterozygosity samples, where HiFi reads are least sortable by parental haplotype and there is more mutual sequence to exploit, but still functions with highly heterozygous F1s. The inter-species GxP trio binned assemblies were more contiguous (+25% NG50) compared to the trio aware assemblies, but contained several large (>5 Mb) misassemblies which the latter did not (Supplementary Figure 1b,c). Phasing in both HiFi- and ONT-based assemblies improved as heterozygosity increased and allowed more cleanly resolved haplotypes, which can be beneficial to downstream analyses (Feng & Li, 2021; Yang & Chaisson, 2021). The ability of hifiasm, and to a lesser extent Shasta with diploid-aware polishing, to assemble phased haplotypes from purebred individuals also avoids ethical and logistical concerns regarding the higher heterozygosity crosses previously targeted (Koren et al., 2018). In situations where parental data is unavailable, accurate haplotype phasing is still possible with supplementary data on the offspring (Cheng, Jarvis, et al., 2021; Porubsky et al., 2020). Our results show that HiFi reads alone are sufficient for contig-level phasing for higher heterozygosity individuals like the GxP (Supplementary Figure 7).

The minigraph pangenomes were strongly comparable whether the input assemblies were all HiFi-based, ONT-based, or a mix. The greater completeness observed in hifiasm assemblies is reflected in those pangenomes containing more non-reference sequence, but no notable HiFi or ONT specific biases were observed in the pangenomes. The quality and completeness of the pangenome backbone can have an impact, seen on the incomplete Angus chromosome 28 or the ARS-UCD1.2 chromosomes generally lacking centromeric sequence, but again we found no specific bias between CLR-or ONT-based backbones. These results indicate that optimal minigraph pangenomes would use high-quality, complete genomes as the backbone, like emerging telomere-to-telomere assemblies (Belser et al., 2021; Nurk et al., 2021; van Rengs et al., 2021). Alternatively, reference-free approaches that use base-level alignment may circumvent these issues and resolve variation down to single nucleotides (Armstrong et al., 2020; Garrison, n.d.), which may prove crucial for breakpoint resolution and related analyses.

Mutual variation identified through shared paths in the pangenome provides opportunities to study the phylogeny of Bovinae beyond SNPs and indels. The ability to accurately separate and represent paths within SVs also enables pangenome-based GWAS (PWAS) (Gupta, 2021), as recently explored in crops (Della Coletta et al., 2021; Hufford et al., 2021; Song et al., 2020). We identified multiple SVs overlapping annotated coding sequences in different bovine pangenomes, demonstrating that pangenomes provide a framework to make them amenable to association mapping. Furthermore, some of these SVs (e.g., tandem duplications in *QRICH2*) are inaccessible from short or noisy long read alignments and some (e.g., an insertion of *HSPA1B*) are challenging to resolve even with long read alignments. These cases highlight the benefits of *de novo*, haplotype-resolved assemblies and pangenome integration which are better able to resolve the variation.

Our work shows that advances in sequencing and algorithms enable computationally feasible haplotype-resolved assembly of 20x HiFi or 60x ONT coverage while retaining 90+% accuracy of detecting SVs when integrated into pangenomes. Structural variation-based pangenomes built from these assemblies demonstrated significant consensus regardless of sequence platform and heterozygosity of the F1. Given the manageable read input needed, it is feasible to produce in the order of several dozens of haplotype-resolved assemblies for specific breeds of cattle. This effort can be de-centralized as no manual curation is needed to produce assemblies of sufficient quality for SV detection, even from purebred individuals. Due to the low effective population size of most breeds of cattle (Stachowicz et al., 2011), the resulting pangenomes would capture the most prevalent SVs within breeds, particularly when assemblies were produced from individuals that account for a large portion of the haplotype diversity of the population (Ros-Freixedes et al., 2017). More haplotype-resolved assemblies are required to reveal rare SVs (Beyter et al., 2021) and characterize SV prevalence in breeds with large effective population size or a history of admixture (J. Kim et al., 2017; K. Kim et al., 2020). Extensive existing short-read sequencing data could then be leveraged to genotype SVs present in the pangenome (Ebler et al., 2020; Hickey et al., 2020) and then impute them into tens of thousands of cattle previously genotyped with microarrays, enabling structural variant analysis at hitherto unattainable scale.

## Methods

### Ethics statement

The sampling of blood from the NxB and OxO trios was approved by the veterinary office of the Canton of Zurich (animal experimentation permit ZH 200/19). All GxP protocols were approved by the Institutional Animal Care and Use Committee (IACUC) of the University of Nebraska–Lincoln, an AAALAC International Accredited institution (IACUC Project ID 1697). Gaur semen was collected and preserved by Omaha’s Henry Doorly Zoo and all protocols were reviewed and approved by their IACUC in 1992.

### Animals

Cows from the Original Braunvieh (O), Brown Swiss (B), and Piedmontese (P) breeds were inseminated with semen samples from Original Braunvieh, Nellore (N), and gaur (G) sires, respectively. The Original Braunvieh sire/dam and Brown Swiss dam were purebred animals with official entries in the respective Swiss herdbooks. The Piedmontese dam was selected by the American Piedmontese Association as representative of their Piedmontese animals. The Nellore sire is from Brazil, with both parents also recorded as purebred Brazilian Nellore. The gaur sire was a zoo animal. Breed affiliation was examined through a PCA on publicly available cattle whole-genome sequencing samples (Supplementary Figure 8).

A female (OxO) and a male (NxB) calf were delivered at term. A female fetus (GxP) cross was collected by cesarean section at 119 days of gestation. Blood samples were taken from the calves and dams by trained veterinarians. Lung tissue was taken from the fetus and DNA extracted as described (Logsdon, 2020). Semen samples were available for the bulls. High-molecular weight DNA was extracted from blood and semen samples, respectively, using Qiagen’s MagAttract HMW DNA Kit as described earlier (Crysnanto et al., 2021).

### Sequencing

The genomes of the F1s were sequenced with long reads using PacBio HiFi and ONT. The PromethION 1D Genomic DNA by ligation SQK-LSK110 library prep kit was used for the OxO, NxB, and GxP F1s. The libraries were respectively sequenced on PromethION (R9.4.1) flowcells (nuclease wash was applied to one cell). The OxO and NxB reads were basecalled on Guppy4, while the GxP reads were basecalled with Guppy3.

Paired-end libraries (2×150 bp) were produced using parental DNA samples and sequenced on Illumina instruments.

### Genome assembly

HiFi reads were first filtered with fastp (version 0.21.0) (S. Chen et al., 2018), removing reads below 1 Kb length or QV20. Nanopore reads were pre-filtered to a minimum of QV7, with no length restrictions. Short reads were also filtered with fastp, using the “-g” parameter to trim polyG tails.

Both HiFi and ONT reads were trio binned into paternal and maternal haplotypes using (Trio)Canu (Koren et al., 2018) (version e0d6bb0), using parental short reads and an estimated genome size of 2.7g.

HiCanu (Nurk et al., 2020) was used with default settings and “genomeSize=2.7g -pacbio-hifi” to produce a draft set of contigs from the HiFi reads. Only contigs which were not suggested as bubbles (suggestBubble=no) were retained, which improved automated coverage detection when using purge_dups (version 1.2.3) (Guan et al., 2020). The final set of “assembled” contigs was obtained after following the default purging pipeline.

The trio aware hifiasm (Cheng, Concepcion, et al., 2021) (version 0.15.3-r339) assemblies were produced from HiFi reads with non-default parameters “-a 5 -n 5”. The resulting graph information was then used in the trio-mode, with parental k-mers constructed with yak (version 0.1-r62-dirty) specified through the “-1 {paternal} -2 {maternal}” parameters. The haplotype gfa files were then converted to fasta contigs with gfa2fa command of gfatools (version 0.5-r234).

The trio binned hifiasm assemblies were obtained with the same assembly parameters, with the addition of the “--primary” flag to produce a primary and alternate assembly. The primary gfa file was converted to fasta as described above. Only the primary assembly was retained.

Peregrine assemblies (Chin & Khalak, 2019) (version main:2aefc14+) were produced using the settings “--with-consensus --shimmer-r 3 --best_n_ovlp 8” on either the trio binned or total HiFi data set where appropriate. Only the primary assembly was retained.

Shasta (Shafin et al., 2020) (version 0.7) assemblies were produced using the standard configuration file “Nanopore-Sep2020.conf”. Assemblies for the Brown Swiss, Nellore, Piedmontese, and gaur haplotypes used trio binned nanopore reads. The OxO had insufficient coverage to trio bin and then assemble, and so the total F1 nanopore read set was used in assembly. Due to the high coverage for the gaur and Piedmontese haplotypes, the parameter “minReadLength=30000” was used instead of the default 10000.

All nanopore assemblies except the Original Braunvieh Shasta were polished using PEPPER (Shafin et al., 2021) (version 0.1) and the trio binned nanopore reads. The Original Braunvieh Shasta assembly was polished with PEPPER-DV (version 0.4), tagging the intermediate bam file with trio binned read IDs to produce the two Original Braunvieh haplotypes. All assemblies were then further polished using short reads. Since short reads cannot be successfully trio binned, we excluded F1 short reads containing e.g. maternal specific k-mers from the short reads used to polish the paternal haplotype and vice versa using meryl. These short reads were then aligned to the long read polished assembly using BWA-mem2 (Li, 2013; Md et al., 2019) and variants were called with DeepVariant (Poplin et al., 2018) (version 1.1). Merfin (Formenti et al., 2021) (commit version 1331fa5) was used to filter the variants and then the assembly was polished using bcftools consensus (Danecek et al., 2021) (version 1.12).

Raven (Vaser & Šikić, 2021) (version 1.5.0) assemblies were obtained from nanopore reads with default parameters, except for “-p 0” to disable racon polishing (Vaser et al., 2017). Assemblies were instead polished with PEPPER for consistency with other ONT assemblers.

Flye (Kolmogorov et al., 2019) (version 2.8.3-b1725) assemblies were constructed with “--genome-size=2.7g --nano-corr” from Ratatosk (version 0.4) (Holley et al., 2021) error-corrected nanopore reads. The nanopore reads were corrected using a reference-guided approach, taking the haplotype-specific hifiasm assembly as the reference. Ambiguous IUPAC codes were randomly replaced with equal probability of an appropriate nucleotide. The contig set was taken from the pre-scaffolding result of Flye and was not polished due to the pre-corrected reads having sufficient accuracy.

### Scaffolding

Contigs were scaffolded into chromosomes by the reference-guided approach of RagTag (Alonge et al., 2021) (v2.0.1) to ARS-UCD1.2, using the additional parameters “--mm2-params “-cx asm5” -r -m 1000000”.

### Assembly merging

Hifiasm and Shasta assemblies were merged using RagTag patch, alternatively using either the hifiasm or Shasta assembly as the base assembly, with the same parameters used for RagTag scaffolding. The merged assembly was then scaffolded to ARS-UD1.2 again as described above.

### Quality metrics

Completeness was assessed with BUSCO (version 5.1.2), using the metaeuk backend (commit 9dee7a7) and odb10 cetartiodactyla database (e96dfc6299c567768085ee9569b6ab15). QV and PG50 were calculated with merqury (version 1.3) (Rhie et al., 2020), with k-mer databases constructed from short reads using meryl (https://github.com/marbl/meryl) (version r953). NG50 was assessed with calN50.js (https://github.com/lh3/calN50). Repetitive elements were identified by RepeatMasker (http://www.repeatmasker.org) (version 4.1.1) and rmblast (version 2.10.0), using a modified version of the 2018 Repbase database. For whole genome analysis, the rush job mode was used, while for pangenome region analysis the slow mode was used.

### Coverage downsampling

Sequencing subsets were made through seqtk (https://github.com/lh3/seqtk) (version 1.3-r115-dirty) with the command “seqtk seq -f {sample}”, including the “-A” flag to drop quality scores where appropriate. In the case of repeat trials, the “-s {seed}” flag was set with a randomly generated 64-bit integer to ensure a unique subsampling of data. The reduced coverage assemblies were conducted identically to the full coverage methods unless explicitly mentioned otherwise.

### Coverage depth estimation

Sequencing reads were mapped against ARS-UCD1.2 with minimap2 (version 2.19-r1059-dirty) (Li, 2018). Coverage depth was determined by megadepth (version 1.1.0c) (Wilks et al., 2021), and averages over 10 Kb windows were estimated with pyBigWig (version 0.3.18) (Ramírez et al., 2016).

### Pangenome construction and bubble extraction

Pangenomes were constructed on a per chromosome basis using minigraph (vesion 0.15-r426) (Li et al., 2020), with default parameters and ARS-UCD1.2 as the initial backbone. The selected assemblies were added to the graph in a randomly shuffled order, as “minigraph -xggs <ARS-UCD1.2> <shuffled input assemblies>“. As input order can affect pangenome construction, this was repeated 20 times with different random orders for each set of inputs (e.g., hifiasm only, Shasta only, mixed, etc.). Pangenomes were visualized with Bandage (version 0.8.1) (Wick et al., 2015).

Haplotype paths were called with minigraph, using the “--call --xasm” option for each respective assembly, resulting in bed files indicating which node the assembly took through each bubble. Intersection sets for the assemblies across each bubble were created with UpSetPlot (https://github.com/jnothman/UpSetPlot) (version 0.6.0). The intersections were then converted into a condensed distance matrix, where each element in the matrix indicates on how many occasions two assemblies took different paths through the same bubble.

### SNP phylogeny

Parental short reads were mapped to the ARS-UCD1.2 reference using BWA-mem2. Variants were called using DeepVariant (version 1.1) with the “WGS” model and merged using GLnexus (version 1.3.1) (Lin et al., 2018). Phylograms were constructed per chromosome with vcf-kit (version 0.2.9) (Cook & Andersen, 2017), using the command “vk phylo tree upgma”.

### Identifying potential trait-associated SVs

All entries in the ARS-UCD1.2 annotation (RefSeq release 106) with a “CDS” description were extracted into a bed file. Bubbles from the respective hifiasm or Shasta pangenomes were also extracted into bed files using the command “gfatools bubble”. The two bed files were then intersected using bedtools (version 2.30.0) (Quinlan & Hall, 2010) with the intersect command and “-wo” to find SVs that impact coding sequence. We then filtered out SVs where haplotype path information through the bubble was not present for all assemblies, which removed 92 and 77 incomplete or ambiguous associations in hifiasm and Shasta pangenomes.

To assess the significance of the SV-gene overlaps, we collected the lengths for the pangenome bubbles per chromosome and then randomly assigned a new location in the uniform interval [1,chromosome length – bubble length]. These randomly permuted bubble locations were then tested for overlaps with CDS regions as described above. We used a one-sided Wilcoxon signed-rank test, assuming the real bubbles would have fewer overlaps with the CDS regions compared to the permuted bubbles.

The set of overlapped genes was compared against the OMIA database (https://www.omia.org/home/)” /www.omia.org/home/)as well as pLI scores in human orthologues (ftp://ftp.broadinstitute.org/pub/ExAC_release/release1/manuscript_data/forweb_cleaned_exac_r03_march16_z_data_pLI.txt.gz) to produce a table containing all genes of interest overlapping pangenome bubbles. Several specific genes were selected for further investigation based on their presence in OMIA or known existing literature describing trait-associated variants.

### Brown Swiss hifiasm localized reassembly

HiFi reads in the area of the *GC* bubble were extracted from a bam file of the Brown Swiss trio binned reads aligned to ARS-UCD1.2 using samtools over the interval 6:86900000-87100000. These reads were then assembled using hifiasm as described for standard hifiasm trio binned assemblies and then were used in a local pangenome analysis where required.

### OxO haplotypes

We used the maternal haplotype of the OxO as the representant assembly for Original Braunvieh (chosen for its more complete X chromosome although it was eventually not used in our analyses) in all analyses where we compared breed/species. This is the case in Figure 4, Figure 5a-c, Figure 6, Figure 7a-e, and the Supplementary material except Figure_SN_1_16 and Figure_SN_1_17. When intra-breed variation for the Original Braunvieh was relevant, we also included the paternal haplotype of the OxO, used in Figure 5d, Figure 7g,f, and the two Supplementary figures listed above.

### Miscellaneous analysis

Assembly, validation, and pangenome workflows have been implemented using Snakemake (Köster et al., 2021), and are available online (https://github.com/AnimalGenomicsETH/bovine-assembly). Nucleotide and protein sequence logos were generated using the R package ggseqlogo (Wagih, 2017).

## Availability of data

HiFi reads for the OxO and NxB F1s are available at the study accession PRJEB42335 under sample accession SAMEA7759028 and SAMEA7765441.

ONT reads for the OxO and NxB are available at the study accession PRJEB42335 under sample accession SAMEA10017983 and SAMEA10017982.

Short reads for the OxO and NxB are available under accession number SAMEA9986200 and SAMEA7589752. Parental short reads are available at SAMEA9986201 & SAMEA9986199 (OxO) and at SAMEA6163185 & SAMEA9533783 (NxB).

Long and short read sequencing data for the GxP trio are available at the study accession PRJEB48481 under secondary accessions SAMEA10563833, SAMEA10563834, and SAMEA10563835.

## Acknowledgements

We thank Dr. Melissa Terranova and Flavio Ferrari (AgroVet-Strickhof) for animal handling and Dr. Sandra Milena Bernal Ulloa for sampling blood. We are thankful for the technical support provided by Dr. Anna Bratus-Neuenschwander from the ETH Zürich technology platform FGCZ (https://fgcz.ch) for sequencing and DNA fragment analysis. The results reported here were made possible with resources provided by the USDA shared compute cluster (Ceres) as part of the ARS SciNet initiative. We thank the USMARC Core Facility staff for outstanding technical assistance. We also thank B. Lee, J. Carlson, K. McClure, H. Clark, H. Sadd, M. Sadd, and B. Shuck for outstanding technical support. We thank the North American Piedmontese Association for their enthusiastic support and assistance in producing gaur interspecies crosses, and Dr. Jason Herrick and Omaha’s Henry Doorly Zoo for the gaur photograph and semen. Mention of trade names or commercial products in this publication is solely for the purpose of providing specific information and does not imply recommendation or endorsement by the U.S. Department of Agriculture. USDA is an equal opportunity provider and employer.

