## Supplementary Material for "Structural variant-based pangenome construction has low sensitivity to variability of haplotype-resolved bovine assemblies"

Supplementary Table 1. Sequencing read details for HiFi, ONT, and short reads (SR). Breeding refers to the type of F1 crossing. Target library sizes and SMRT cells are reported for HiFi reads, with multiple libraries separated by commas, as well as the total number of bases, reads, and read N50. ONT data did not have a target library size, but the number of PromethION cells and nuclease washes are reported along with reads, bases, and read N50. Short reads for both the F1s and sires/dams libraries are 150 bp in length, with the number of bases reported.

| Sire/dam bases | 83974454313/ 90279569069 | 145456304453/ 152857605524 | 97052058593/122689575398 |
| --- | --- | --- | --- |
| F1 SR bases | 83420372505 | 61158863267 | 99681429463 |
| ONT N50 | 63993 | 44830 | 48583 |
| ONT bases/reads | 97741535249/ 2617673 | 277704422221/ 9066711 | 410851293648/ 15752473 |
| ONT cells (nuclease washes) | 2 (0) | 3 (0) | 16 (1) |
| HiFi N50 | 19621 | 20792 | 12914 |
| HiFi bases/reads | 86877013947/ 4491846 | 140232393384/ 6803270 | 139087513200/ 9359887 |
| HiFi library size in kb (cells) | 18.7 (4) | 21 (5), 15 (1) | 10 (9), 15 (4) |
| F1 Sex | Female | Male | Female |
| Breeding | Within-breed | Inter-subspecies | Inter-species |
| F1 | OxO | NxB | GxP |

Supplementary Table 2. Trio binning of HiFi and ONT long read data. For each F1, the number of heterozygous sites is estimated by aligning short reads to ARS-UCD1.2, calling variants, and counting the number of heterozygous sites. Domintette, the cow on which ARS-UCD1.2 is based, had 2,607,461 heterozygous sites. The percentage of bases/reads assigned to each haplotype, or an unknown bin, are given for each trio. Separability refers to the percentage of F1 bases assigned to either haplotype 1 or 2.

|  | | HiFi | | | | ONT | | | |
| --- | --- | --- | --- | --- | --- | --- | --- | --- | --- |
| F1 | Heterozygous sites | Hap1 | Hap2 | Unknown | Separability | Hap1 | Hap2 | Unknown | Separability |
| OxO | 4,024,944 | 41.5/40.7 | 39.6/38.8 | 18.9/20.5 | 81.12 | 51.4/49.8 | 48.2/47.0 | 0.4/3.2 | 99.67 |
| NxB | 11,506,532 | 48.1/48.2 | 50.7/50.8 | 1.0/1.2 | 98.99 | 46.1/46.7 | 53.9/52.9 | 0.0/0.4 | 99.97 |
| GxP | 23,190,285 | 49.0/49.0 | 50.9/50.9 | 0.1/0.1 | 99.88 | 47.0/43.4 | 52.9/48.9 | 0.1/7.7 | 99.83 |

Supplementary Table 3. Assembly metrics for the additional four assemblers. Columns as are described in Table 1. Due to lower HiFi and ONT coverage for the Original Braunvieh, primary assemblies (i.e., not haplotype-resolved) were assembled where necessary and are marked by *. Assemblies are available online (Leonard, 2022).

| Breed or species | Read | Assembler | Size (autosomal size) | Contigs (autosomal contigs) | NG50 | | PG50 | QV | BUSCO (single copy) | | Repeat | |
| --- | --- | --- | --- | --- | --- | --- | --- | --- | --- | --- | --- | --- |
| Original Braunvieh | HiFi | Peregrine* | 3.34 (2.56) | 3108 (134) | 43.6 | | 1.02 | 48.0 | 95.7 (93.6) | | 51.57 | |
|  |  | HiCanu* | 3.22 (2.56) | 1729 (116) | 56.4 | | 1.51 | 51.1 | 96.2 (94.2) | | 50.63 | |
|  |  | HiCanu | 3.03 (2.54) | 3240 (822) | 7.36 | | 7.22 | 48.7 | 95.3 (93.4) | | 48.01 | |
|  | ONT | Raven* | 2.70 (2.52) | 459 (283) | 20.0 | | 0.32 | 37.1 | 87.3 (85.8) | | 42.75 | |
|  |  | Flye* | 2.66 (2.50) | 1344 (267) | 60.8 | | 0.36 | 42.8 | 95.9 (94.2) | | 42.04 | |
| Nellore | HiFi | HiCanu | 2.85 (2.52) | 3266 (153) | 39.2 | | 32.8 | 44.5 | 93.4 (91.7) | | 47.37 | |
|  |  | Peregrine | 2.91 (2.56) | 1302 (127) | 38.7 | | 38.3 | 45.9 | 92.9 (91.4) | | 47.31 | |
|  | ONT | Raven | 2.52 (2.50) | 185 (131) | 62.4 | | 22.3 | 33.3 | 86.5 (85.2) | | 41.43 | |
|  |  | Flye | 2.53 (2.50) | 585 (130) | 71.5 | | 25.7 | 43.0 | 93.1 (91.6) | | 41.71 | |
| Brown Swiss | HiFi | HiCanu | 3.10 (2.51) | 3969 (182) | 29.0 | | 27.3 | 43.6 | 96.1 (94.2) | | 49.67 | |
|  |  | peregrine | 3.09 (2.54) | 1805 (206) | 25.6 | | 22.2 | 44.9 | 95.8 (94.1) | | 49.11 | |
|  | ONT | Raven | 2.74 (2.50) | 366 (132) | 54.3 | | 36.7 | 32.3 | 95.5 (93.9) | | 43.71 | |
|  |  | Flye | 2.65 (2.49) | 553 (111) | 68.2 | | 12.7 | 42.5 | 95.7 (94.1) | | 42.12 | |
| gaur | HiFi | HiCanu | 2.91 (2.53) | 1099 (119) | 41.1 | | 39.1 | 50.3 | 95.7 (93.9) | | 46.21 | |
|  |  | Peregrine | 2.94 (2.52) | 2402 (679) | 5.7 | | 5.4 | 46.5 | 94.8 (93.0) | | 46.73 | |
|  | ONT | Raven | 2.65 (2.49) | 212 (11) | 65.8 | | 62.2 | 36.8 | 94.8 (93.0) | | 41.96 | |
|  |  | Flye | 2.62 (2.48) | 345 (145) | 68.4 | | 68.2 | 42.1 | 93.4 (91.6) | | 41.86 | |
| Piemontese | HiFi | HiCanu | 3.03 (2.53) | 1575 (119) | 47.9 | | 41.9 | 48.9 | 95.9 (94.2) | | 47.81 | |
|  |  | Peregrine | 3.04 (2.52) | 2886 (771) | 5.2 | | 4.7 | 46.5 | 94.7 (93.0) | | 47.71 | |
|  | ONT | Raven | 2.74 (2.50) | 414 (140) | 61.6 | | 61.3 | 31.9 | 94.9 (93.2) | | 43.57 | |
|  |  | Flye | 2.64 (2.49) | 325 (98) | 70.0 | | 69.9 | 44.1 | 95.7 (94.1) | | 41.95 | |
| Hereford (ARS-UCD1.2) | CLR | Canu | 2.72 (2.49) | 2597 (289) | 25.9 | | N/A | 35.8 | 95.7 (93.9) | | 42.96 | |
| VGP Standards | | | | | | 1 | 0.1 | 40 | | 90 | | N/A |

Supplementary Table 4. Repetitive content of sequence that was not assigned to chromosomes. Contigs which were scaffolded to unplaced sequence in ARS-UCD1.2 (typically denoted NKL…) are labelled “REF”, while contigs which did not align confidently to any existing reference sequence are labelled “ASM”. The total number of bases in each contig type are shown in megabases, along with the percentage of repetitive bases identified by RepeatMasker for each major classes, as well as the total percentage of masked bases. The average masked percentage is higher for the 10 produced assemblies compared to ARS-UCD1.2, as expected given the higher potential of HiFi and ONT reads to assemble repetitive regions.

| Breed/Species | Assembler | Tigs | Bases (Mb) | SINEs | LINEs | LTR | Satellites | Unclassified | Total masked |
| --- | --- | --- | --- | --- | --- | --- | --- | --- | --- |
| Hereford (ARSUCD1.2) | Canu | REF | 87.438 | 2.5 | 7.9 | 27.1 | 51.4 | 6.7 | 73.7 |
| gaur | hifiasm | REF | 324.624 | 0.6 | 0.3 | 16.2 | 79.4 | 3.1 | 89.9 |
|  |  | ASM | 22.850 | 1.1 | 1.9 | 18.5 | 77.3 | 3.7 | 90.0 |
|  | Shasta | REF | 8.168 | 0.4 | 0.9 | 20.1 | 79.3 | 2.3 | 91.4 |
|  |  | ASM | 8.336 | 3.6 | 3.5 | 47.3 | 32.2 | 11.1 | 70.0 |
| Piemontese | hifiasm | REF | 295.895 | 1.1 | 0.7 | 22.4 | 71.4 | 5.1 | 86.5 |
|  |  | ASM | 97.389 | 1.8 | 5.3 | 28.4 | 58.6 | 4.9 | 78.9 |
|  | Shasta | REF | 16.705 | 0.9 | 3.5 | 25.6 | 73.9 | 4.2 | 91.1 |
|  |  | ASM | 17.535 | 1.8 | 5.1 | 22.7 | 68.7 | 5.3 | 86.2 |
| Nellore | hifiasm | REF | 232.134 | 1.0 | 0.5 | 24.4 | 74.1 | 5.1 | 90.0 |
|  |  | ASM | 50.400 | 0.9 | 2.1 | 25.8 | 71.7 | 3.5 | 88.2 |
|  | Shasta | REF | 18.208 | 0.4 | 0.9 | 29.6 | 74.2 | 2.1 | 90.7 |
|  |  | ASM | 31.937 | 1.1 | 3.8 | 19.5 | 74.0 | 3.8 | 88.1 |
| Brown Swiss | hifiasm | REF | 238.508 | 1.3 | 2.7 | 22.2 | 66.4 | 5.5 | 82.8 |
|  |  | ASM | 70.120 | 0.5 | 1.0 | 21.5 | 80.1 | 2.0 | 92.8 |
|  | Shasta | REF | 43.099 | 0.7 | 4.2 | 22.1 | 71.3 | 3.8 | 86.9 |
|  |  | ASM | 22.063 | 1.5 | 4.9 | 21.1 | 70.3 | 4.2 | 86.2 |
| Original  Braunvieh | hifiasm | REF | 307.811 | 1.0 | 0.5 | 25.6 | 74.0 | 4.5 | 90.2 |
|  |  | ASM | 79.682 | 1.4 | 3.7 | 28.3 | 67.0 | 4.4 | 86.1 |
|  | Shasta | REF | 33.545 | 0.6 | 1.6 | 34.1 | 70.0 | 3.6 | 90.4 |
|  |  | ASM | 43.033 | 1.3 | 4.4 | 25.7 | 69.9 | 5.0 | 88.2 |

Supplementary Table 5. Statistics for hifiasm and Shasta pangenomes. N_b_ is the total number of bubbles identified in the pangenome. L_ref_ and L_b_ refer to the mean length of the reference path and non-reference paths through bubbles. N_Ins_ and N_Del_ are respectively the number of predicted insertion or deletion events, for when the reference path is of length zero and non-reference path has non-zero length, or the reverse. K is the mean number of non-reference nodes in bubbles.

|  | N_b_ | L_ref_ | L_b_ | N_Ins_ | N_Del_ | K |
| --- | --- | --- | --- | --- | --- | --- |
| hifiasm | 98,054 | 594 | 1080 | 27,642 | 24,456 | 1.755 |
| Shasta | 98,545 | 584 | 924 | 27,512 | 24,294 | 1.762 |

Supplementary Table 6. Pangenome bubbles unique to HiFi or ONT assemblies. Values are averaged across 10 pangenomes constructed from five hifiasm and five Shasta assemblies. Bubbles are considered to be unique to HiFi or ONT if they are 1+ Kb in size and not within 1 Kb of a bubble from the other read technology. Bases are listed in Mb, with the repeat types given as percentages.

| Read | Bubbles | Bases (Mb) | SINEs | LINEs | LTR | Satellites | Unclassified | Total Masked |
| --- | --- | --- | --- | --- | --- | --- | --- | --- |
| HiFi | 57 | 1.85 | 5.9 | 22.9 | 5.8 | 4.3 | 13.0 | 54.0 |
| ONT | 36 | 0.39 | 8.4 | 12.9 | 4.0 | 3.8 | 14.9 | 45.3 |

Supplementary Table 7. Quality metrics for downsampled assemblies. Coverage for hifiasm assemblies is diploid (denoted by *), while Shasta coverage is the trio binned coverage for the respective haplotype (except for Original Braunvieh, which is diploid coverage followed by polishing-based phasing).

| Breed/species | Read technology | Coverage | Size (autosomal size) | Contigs (autosomal contigs) | NG50 | PG50 | QV | BUSCO (single copy) | Repeat |
| --- | --- | --- | --- | --- | --- | --- | --- | --- | --- |
| Original  Braunvieh | hifiasm | 19.3* | 2.96 (2.53) | 3266 (1654) | 2.9 | 2.3 | 43.5 | 92.7 (90.3) | 46.5 |
|  | Shasta | 29.0 | 2.70 (2.48) | 2104 (114) | 50.6 | 2.3 | 39.8 | 95.1 (93.4) | 43.2 |
| Brown Swiss | hifiasm | 20.3* | 2.92 (2.53) | 3015 (1789) | 2.3 | 2.2 | 40.7 | 93.7 (92.1) | 46.4 |
|  | Shasta | 24.4 | 2.68 (2.48) | 2456 (418) | 10.4 | 10.0 | 40.0 | 94.7 (93.1) | 43.0 |
| Nellore | hifiasm | 20.3* | 2.81 (2.54) | 3016 (1863) | 2.2 | 2.2 | 40.9 | 91.1 (89.6) | 45.9 |
|  | Shasta | 20.9 | 2.55 (2.48) | 2399 (399) | 10.9 | 10.8 | 40.1 | 92.5 (91.0) | 42.8 |
| gaur | hifiasm | 24.2* | 2.90 (2.50) | 4768 (2055) | 2.0 | 1.8 | 42.4 | 94.1 (92.3) | 45.8 |
|  | Shasta | 23.6 | 2.64 (2.48) | 1065 (411) | 13.5 | 13.5 | 38.4 | 94.1 (92.3) | 42.0 |
| Pied | hifiasm | 24.2* | 2.97 (2.51) | 5176 (2128) | 1.9 | 1.7 | 42.3 | 94.0 (92.3) | 46.5 |
|  | Shasta | 26.5 | 2.63 (2.47) | 1459 (419) | 11.6 | 11.6 | 37.9 | 94.3 (92.3) | 42.2 |

Supplementary Table 8. Full set of genes overlapped by hifiasm or Shasta pangenome SVs. ARS-UCD1.2 coordinates for the start and end of the overlapped coding sequence region are given for both hifiasm (H) and Shasta (S) pangenomes, along with the number of coding sequence regions overlapped, the mean length of the pangenome bubbles, and the mean number of non-reference paths through the bubble. OMIA indicates if the gene is present in the OMIA database and pLI is the pLI score of the gene (0 if not available).

<CSV FILE>

Supplementary Table 9. NG50 for merged hifiasm and Shasta assemblies. Assemblies were merged with RagTag for each assembly pair separately, either merging the Shasta assembly into the hifiasm assembly (Merge [HiFi-backed]) or vice versa. Contig NG50 values are given in Mb.

|  | Breeds/species | | | | | |
| --- | --- | --- | --- | --- | --- | --- |
|  | O (pat) | O (mat) | N | B | G | P |
| HiFi | 56.0 | 47.0 | 94.4 | 86.7 | 73.5 | 52.0 |
| ONT | 71.6 | 71.7 | 68.5 | 64.0 | 68.1 | 82.8 |
| Merge (HiFi-backed) | 62.4 | 65.6 | 96.4 | 86.7 | 73.5 | 65.2 |
| Merge (ONT-backed) | 72.9 | 72.9 | 68.5 | 64.0 | 73.7 | 84.1 |

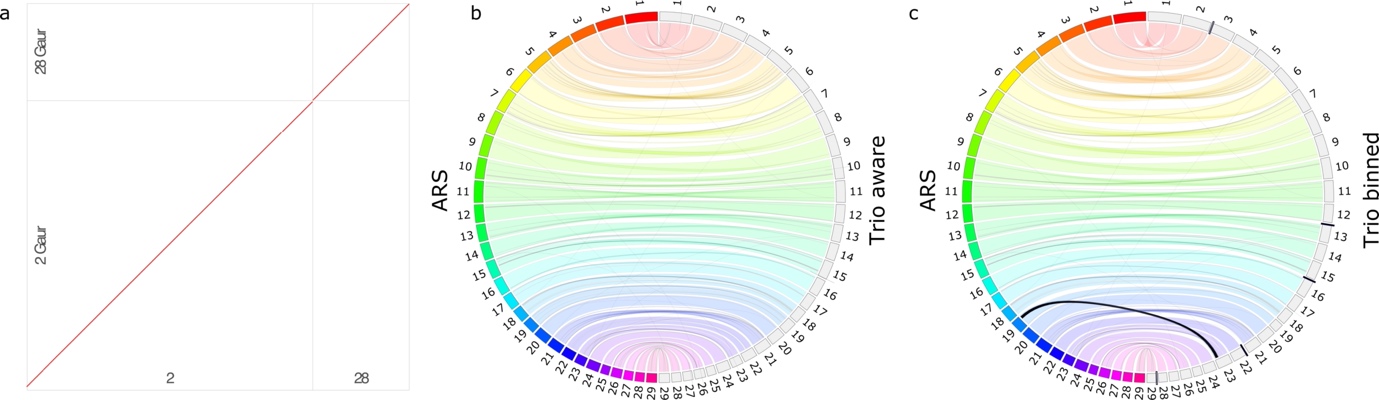

Supplementary Figure 1. Alignment plots of hifiasm gaur assemblies and ARS-UCD1.2. a) Dot plot of chromosomes 2 and 28 in gaur (left) and ARS-UCD1.2 (bottom). Since the gaur assembly did not span the centromeric fusion, the scaffolding to chromosomes 2 and 28 behave normally. b) The trio aware assembly of the GxP gaur haplotype has no major misassemblies compared to ARS-UCD1.2, while the trio binned assembly in c) has several, like 20 Mb of sequence from chromosome 19 is assembled into chromosome 23. Similarly, the trio binned assembly has greater standard deviation in autosome size compared to ARS-UCD1.2 than the trio aware assembly (± 6.2 versus 1.4 Mb). After accounting for differences in centromeric sequence, the trio binned gaur assembly had more than 5 Mb of sequence potentially missing on chromosomes 3 and 29, while it also had more than 5 Mb of potentially spurious sequence on chromosomes 13, 16, and 22. The trio binned assembly does have more autosomal centromeric sequence (44 versus 17 Mb) but appears to be missing nearly 23 Mb of non-centromeric sequence (9 Mb of which is repetitive). This is reflected in the higher BUSCO score for the trio aware mode compared to trio binned (95.7% versus 94.2% complete BUSCOs).

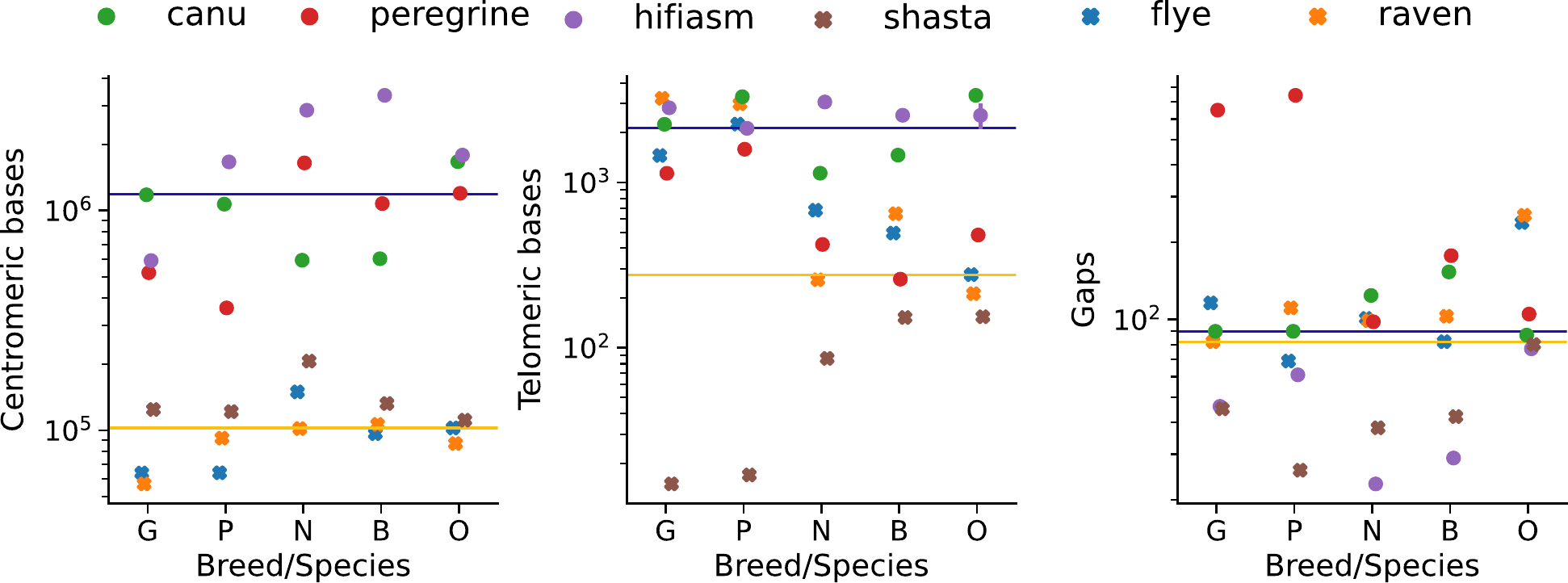

Supplementary Figure 2. Centromere and telomere completeness. HiFi-based assemblies (circles) generally outperform ONT-based assemblies (crosses) for mean centromere and telomere completeness across the autosomes for the different breeed/species assembled. However, ONT-based assemblies have slightly fewer gaps. Error bars represent the 95% confidence interval. Mean values across each sequencing technology are marked by blue (HiFi) and orange (ONT) lines. Shasta tends to under-perform relative to the ONT average for telomere completeness, potentially due to the marker-graph assembly strategy, while hifiasm tends to over-perform relative to HiFi average for the number of autosome gaps.

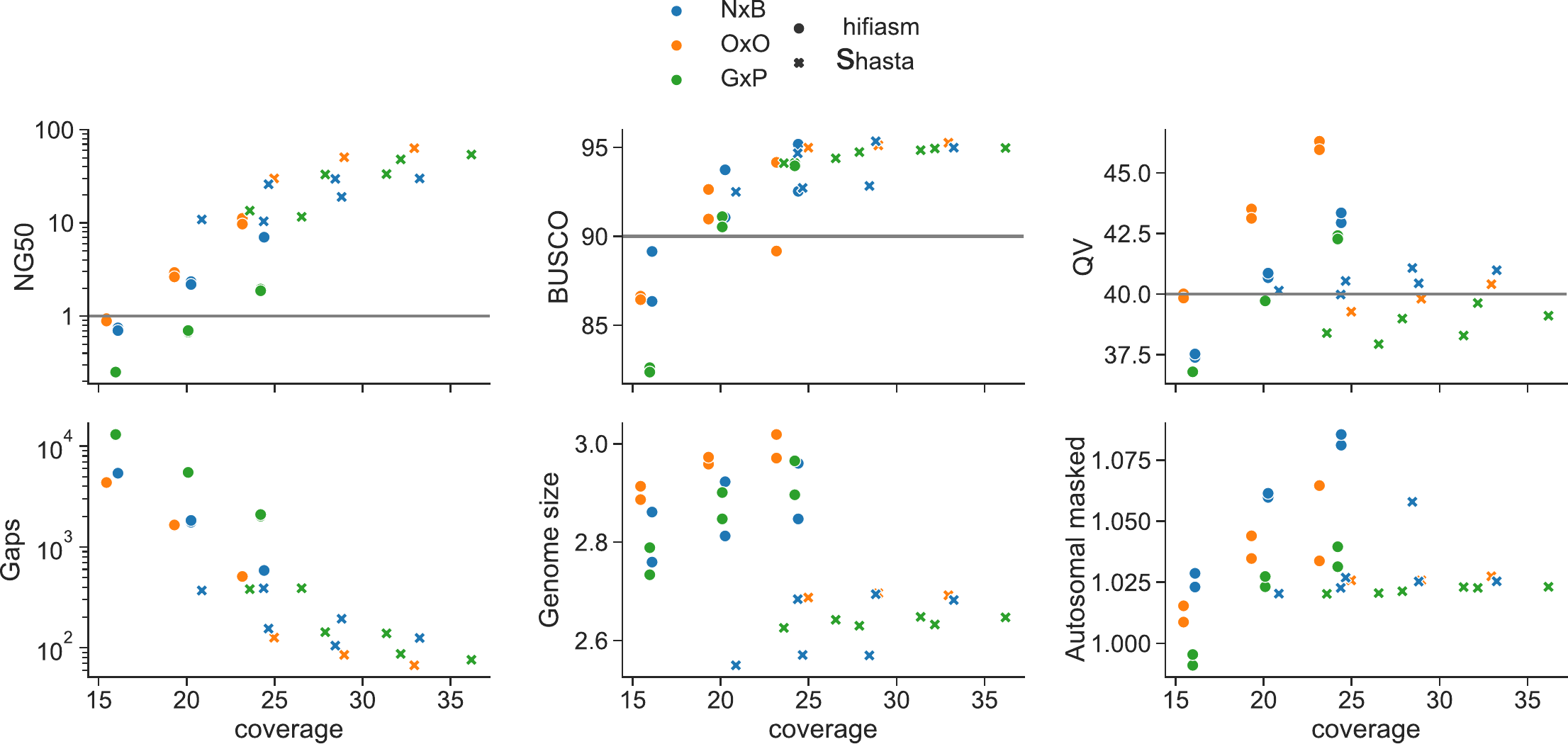

Supplementary Figure 3. Assembly quality metrics for “medium” coverage. The metrics are the same as described in Figure 3. Results are overall similar between the three F1s, where approximately 20x diploid HiFi coverage or 30x haploid ONT coverage is sufficient to produce assemblies at or above the VGP quality standards.

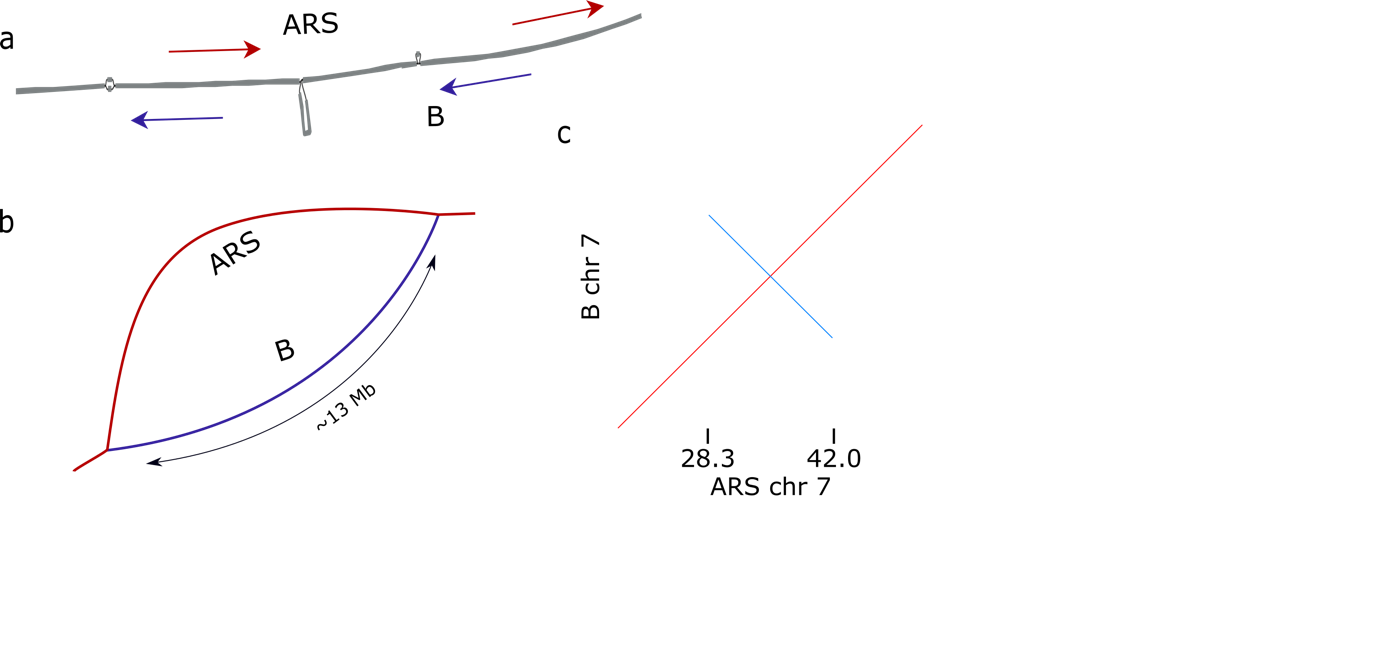

Supplementary Figure 4. Palindromic inversion on BTA7. Minigraph either maps chromosome 7 of the hifiasm assembly of the Brown Swiss haplotype as a) the negative strand (blue arrows) of many small existing nodes in the graph (red arrows) or b) one large, inverted bubble. The latter case generally happens when the Brown Swiss hifiasm assembly is added first. c) A dot-plot of this region aligned between ARS-UCD1.2 and the Brown Swiss hifiasm assembly indicate the region between BTA7:28.3-42.0 Mb is palindromic with support for forward (red) and reverse (blue) alignments. The alignment score is only slightly higher for the reverse alignment compared to the forward, corresponding to graph cases a) and b) respectively.

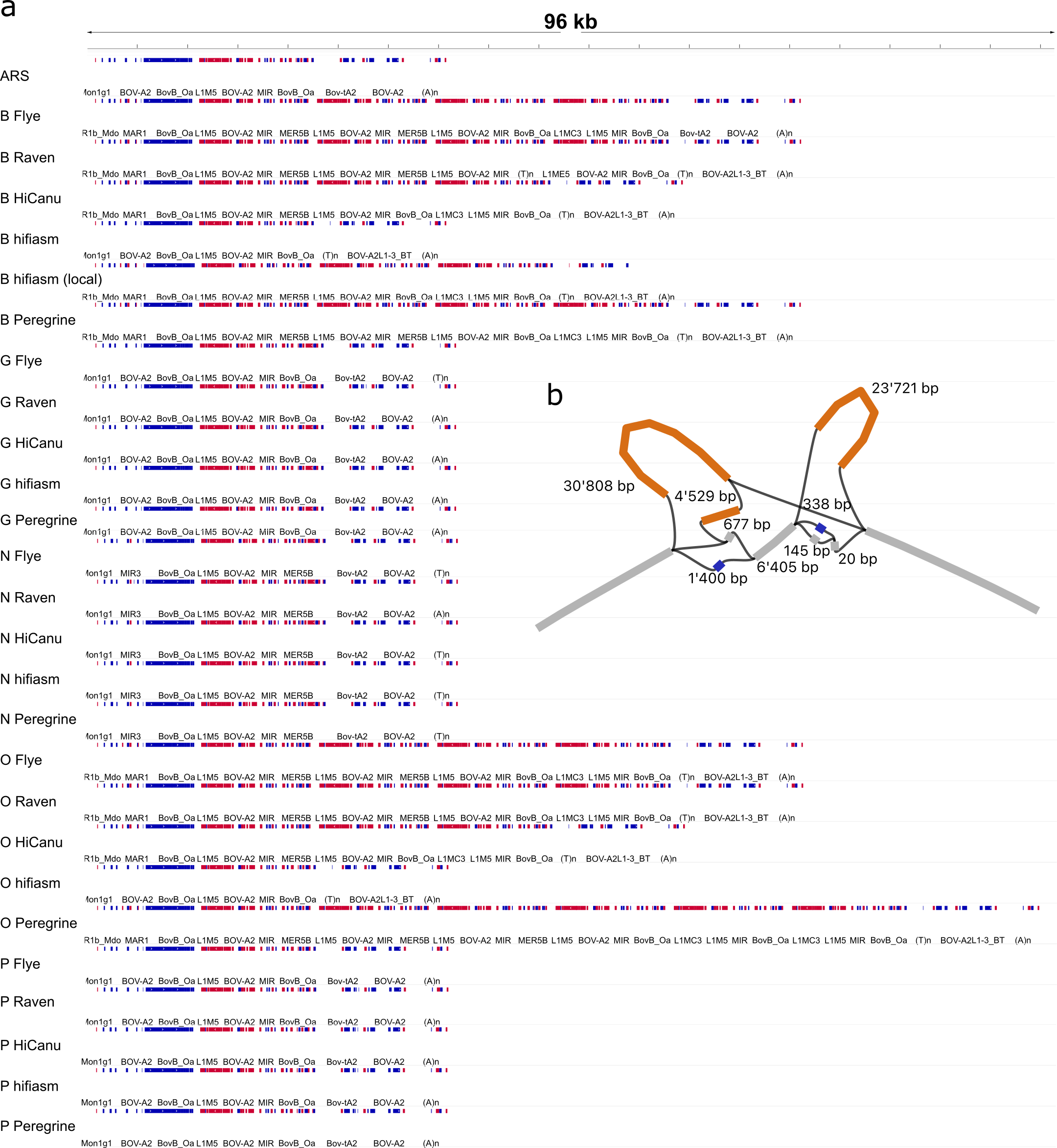

Supplementary Figure 5. GC SV copy number is highly inconsistent across assemblers. a) The same repeat analysis as described in Figure 6 confirms the tandem duplication is never encountered in gaur (G), Nellore (N), and Piemontese (P) across all examined assemblers. However, the duplication copy number for Original Braunvieh ranges from zero to five, while Brown Swiss (B) ranges from zero to three copies. A local re-assembly of the relevant Brown Swiss HiFi reads (align reads to ARS-UCD1.2, and then extract all reads in a ± 20 Kb interval around the ARS-UCD1.2 coordinates of the SV) with hifiasm produces two copies, confirming the ability to resolve this CNV is not a limitation of hifiasm but potentially a sensitivity issue when assembling the whole genome. b) The 31 assembly pangenome still identifies the CNV as well as the 200 bp and 700 bp insertions in gaur and Nellore.

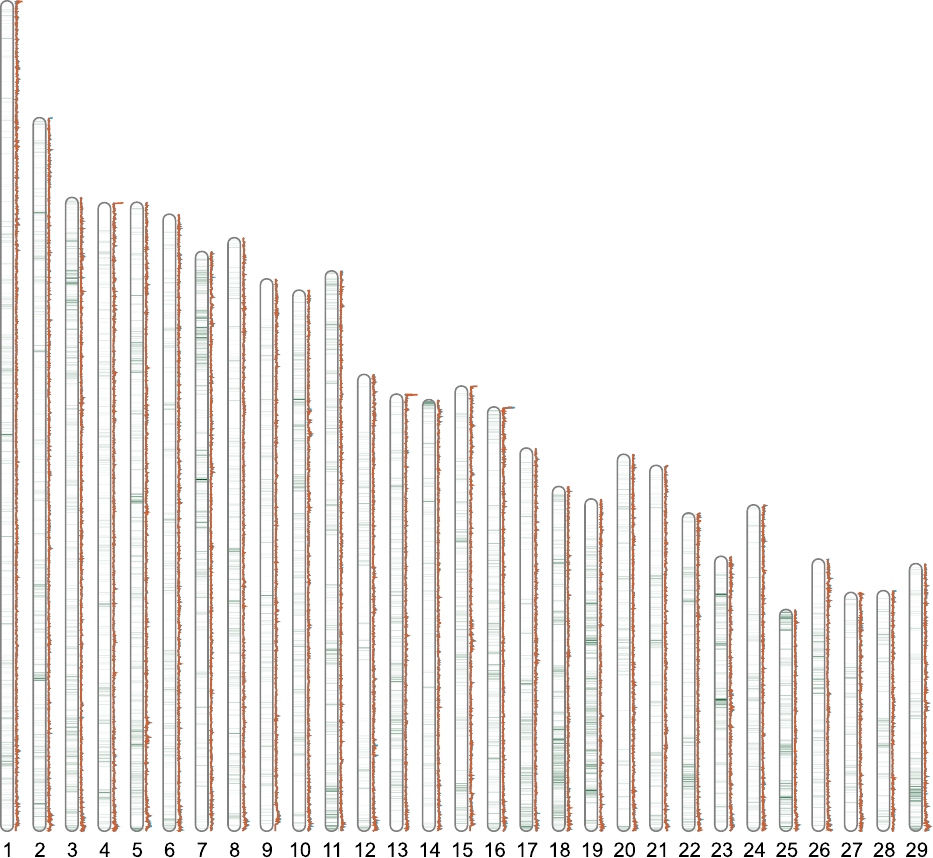

Supplementary Figure 6. Overlaps between protein coding regions and pangenome bubbles. The number of CDS regions in the ARS-UCD1.2 RefSeq 106 annotation per 100 Kb window are shown as a heatmap (white – low to green – high) along the autosomes. Similarly, the number of bubbles from HiFi (blue) and ONT (orange) pangenomes are shown on the right of each autosome. There are some fluctuations between the two bubble counts, particularly at chromosome ends, but there are 3.96 and 3.98 bubbles per 100 Kb window for HiFi and ONT respectively. The root mean square difference between them is 0.93.

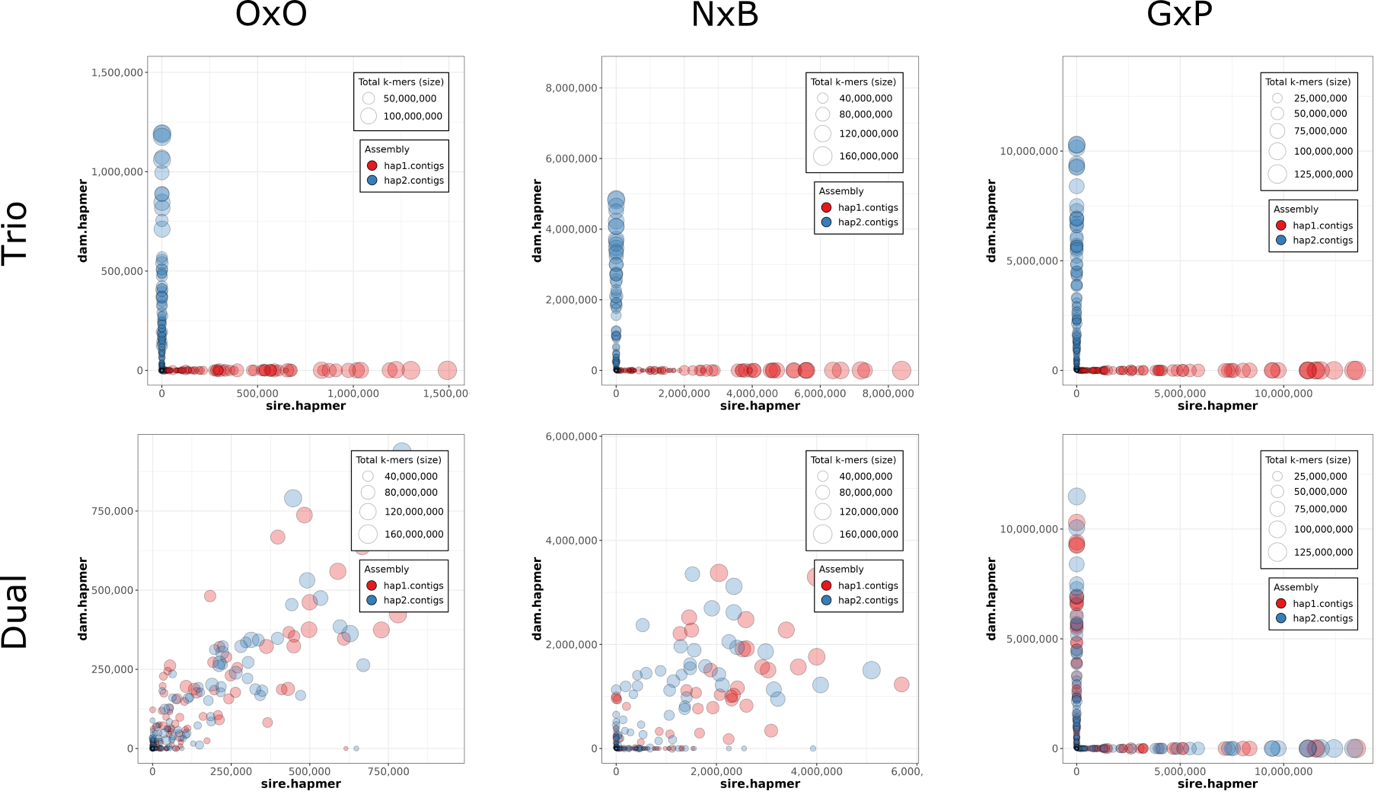

Supplementary Figure 7. Merqury phasing blob plots for OxO, NxB, and GxP hifiasm assemblies using parental information (trio) or using a local phasing algorithm in hifiasm using only the F1 HiFi reads (dual). Blobs represent individual contigs, and the axes represent how many specific dam/sire haplotype k-mers (hapmers) the contig contained. Hap1 and hap2 refers to the paternal and maternal haplotypes respectively. Phasing is highly accurate when using parental information (top). Without parental information, increasing heterozygosity allows improved phasing. The mean shortest Euclidian distance for each blob to the diagonal (greater distance indicates a haplotype is more strongly maternal or paternal) increases for the OxO, NxB, and GxP from 3204.5 to 36322.6 and then 138001.4. Hifiasm cannot infer if a contig belongs to the maternal or paternal assembly since there is no parental information but, given enough heterozygosity (e.g., the GxP), is able to consistently phase haplotypes to the contig level.

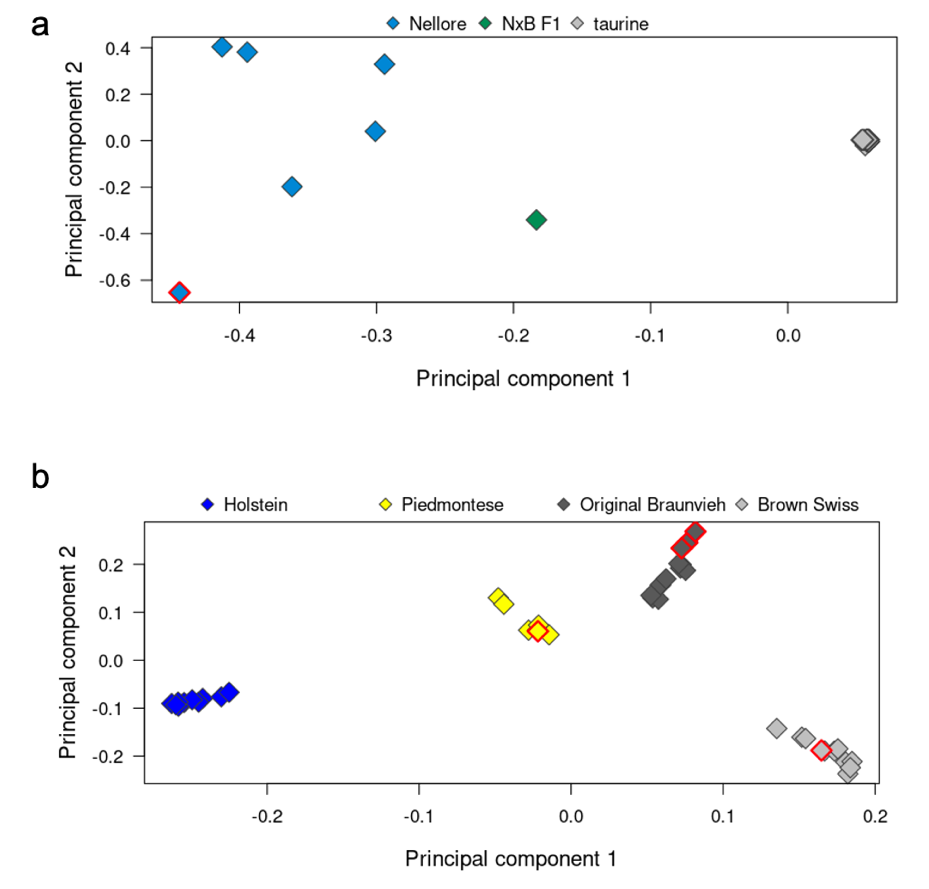

Supplementary Figure 8. PCA of taurine (Holstein, Piedmontese, Brown Swiss, Original Braunvieh), indicine (Nellore) and indicine x taurine (NxB) crosses. Different colors separate individuals by breed. Red framed symbols highlight the purebred animals from the three trios. The PCA was performed for all samples (a) and for the taurine-only (N=41) samples (b). Multi-sample variant calling was done on publicly available whole-genome sequencing data of 10 Brown Swiss (SAMEA5415485, SAMEA7573645, SAMEA7573582, SAMEA7573592,SAMEA7573599, SAMEA5714971, SAMEA6163191, SAMEA7573585, SAMEA6163179, SAMEA6163195), 10 Original Braunvieh (SAMEA4827668, SAMEA5059753, SAMEA5159850, SAMEA6272091, SAMEA4827645, SAMEA4827672, SAMEA6272103, SAMEA6272093, SAMEA5564728, SAMEA5059754), 10 Holstein (SAMEA6528904, SAMEA6528905, SAMEA6528906, SAMEA6528907, SAMEA6528908, SAMEA6528909, SAMEA7015110, SAMEA7015112, SAMEA7015113, SAMEA7015115), 6 Piedmontese (SAMEA5159836, SAMEA19324918, SAMEA33001918, SAMN02941216, SAMN02941219, SAMN02941218) and 5 Nellore (SAMN10486398, SAMN10486400, SAMN10486401, SAMN10486399, SAMN03387027) samples together with short sequencing reads from the OxO and NxB trio as well as the dam of the GxP trio. There were 35.4 million and 20.1 million polymorphic sites along the autosomes used to construct a genomic relationship matrix for the taurine/indicine and taurine-only breeds, respectively.

Supplementary Note 1: Verification of structural variants discovered from the pangenome

After identifying pangenome bubbles overlapping annotated coding sequences, we verified multiple examples by inspecting alignments of long and short sequencing reads against the ARS-UCD1.2 reference sequence. Trio binning and alignment against ARS-UCD1.2 are as described in the Methods. To verify bubbles from the pangenome, we inspected the read alignments at the corresponding positions with the Integrative Genomics Viewer (IGV, (Robinson et al., 2011)).

**Recovery of a tandem duplication upstream the group-specific component (*GC*) gene**

Alignments of long reads confirmed the presence of a large tandem duplication in the promoter region of GC (ENSBTAG00000013718) in the Brown Swiss and Original Braunvieh haplotype (Figure SN1_1). A sequence of 11,780 bp (between 86,949,653 bp and 86,961,433 bp) is duplicated and occurs as between 3 and 4 copies in these haplotypes. This duplication is not present in the gaur, Nellore and Piedmontese haplotype. However, the pangenome revealed two insertions of approximately 178 bp and 730 bp in both the Nellore and gaur haplotypes, that coincide with the tandem duplication (Figure SN_1_2).

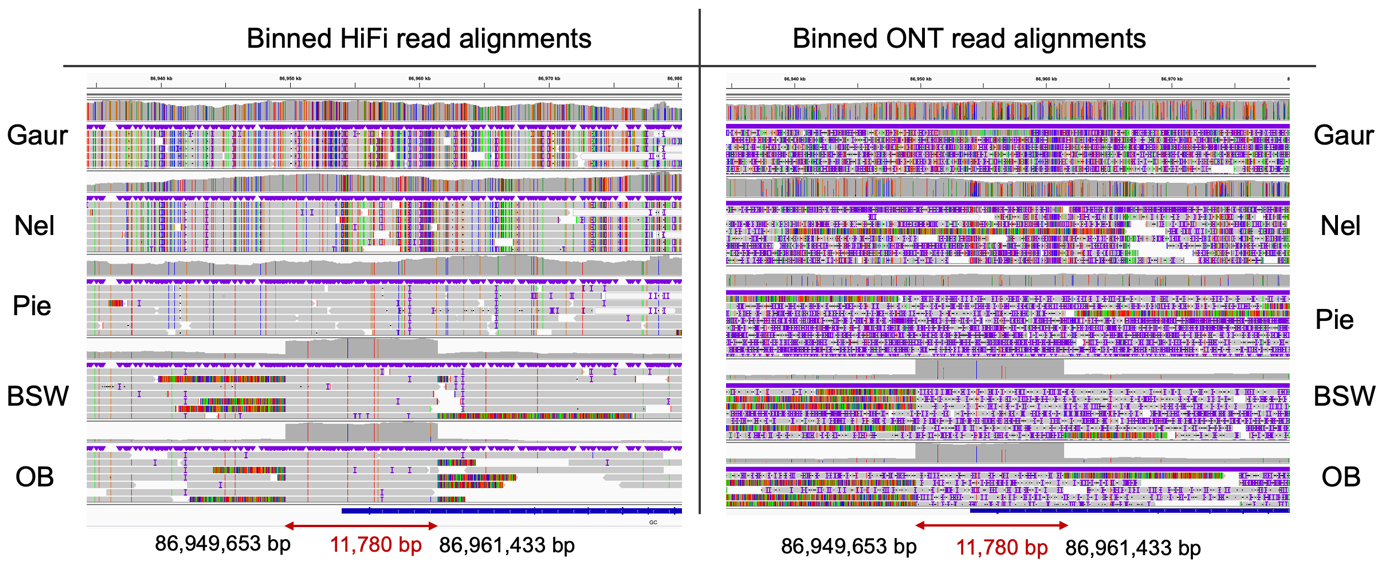

Figure SN1_1: Binned HiFi (left) and ONT (right) read alignments against ARS-UCD1.2. The tandem duplication is visible as elevated coverage in BSW and OB between 86,949,653 bp and 86,961,433 bp.

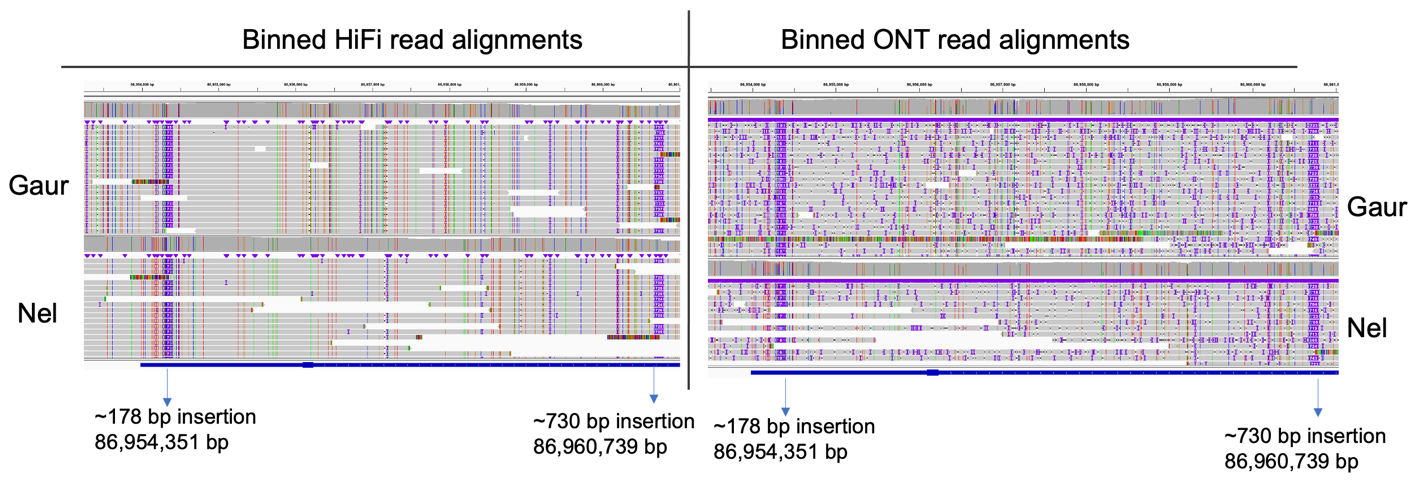

Figure SN_1_2: Binned HiFi (left) and ONT (right) read alignments against ARS-UCD1.2 indicate two insertions of 178 bp and 730 bp in the gaur and Nellore assemblies.

**Structural variation in the promoter region of *ASIP* encoding agouti signaling protein**

The coat of the NxB was light at birth but turned almost black with age (Figure SN1_3). This darkening with age is frequently observed in Brown Swiss, Original Braunvieh, and Nellore cattle. However, the genetic determinants are largely unknown. A recent study discovered a structural rearrangement in the promoter of *ASIP* (ENSBTAG00000034077) to be associated with variation in coat color in Nellore cattle (Trigo et al., 2021). The analysis of short read alignments indicated that the Nellore sire indeed is a heterozygous carrier of the 1155 bp deletion and 150 bp insertion of Bov-tA SINE at CHR13:63,599,803–63,600,957 that is associated with darkness of coat (Trigo et al., 2021). This structural rearrangement is not present in the NxB suggesting it inherited the haplotype without the deletion/insertion from its sire.

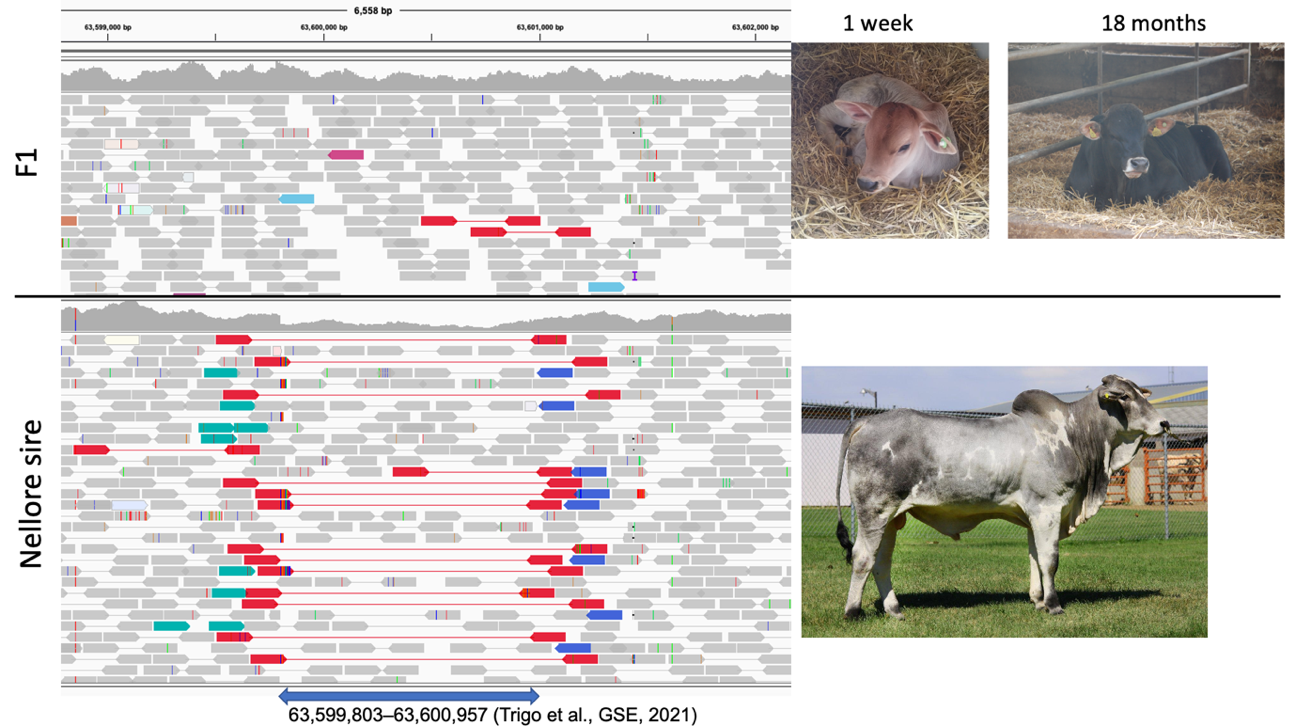

Figure SN1_3: IGV screenshots and photographs from the NxB (upper panel) and its Nellore sire (lower panel). Different colors in the IGV screenshot of the Nellore sire represent reads with abnormal insert size indicating the 1155 bp deletion and 150 bp insertion.

We observed extended segments of elevated and reduced coverage in the promoter region of *ASIP* in the haplotype-resolved read alignments (Figure SN_1_4). Deletion of 8,403 bp sequence between 63,639,803 bp and 63,648,206 bp was visible in all assemblies. In Brown Swiss, this deletion partly overlapped with a 3,288 bp region of elevated coverage between 63,644,918 bp and 63,648,206 bp. Read coverage varied greatly across the assemblies at a 1278 bp region between 63,629,629 bp and 63,630,907 bp. However, an immediate association between any of the detected large structural variants and coat color was not readily apparent.

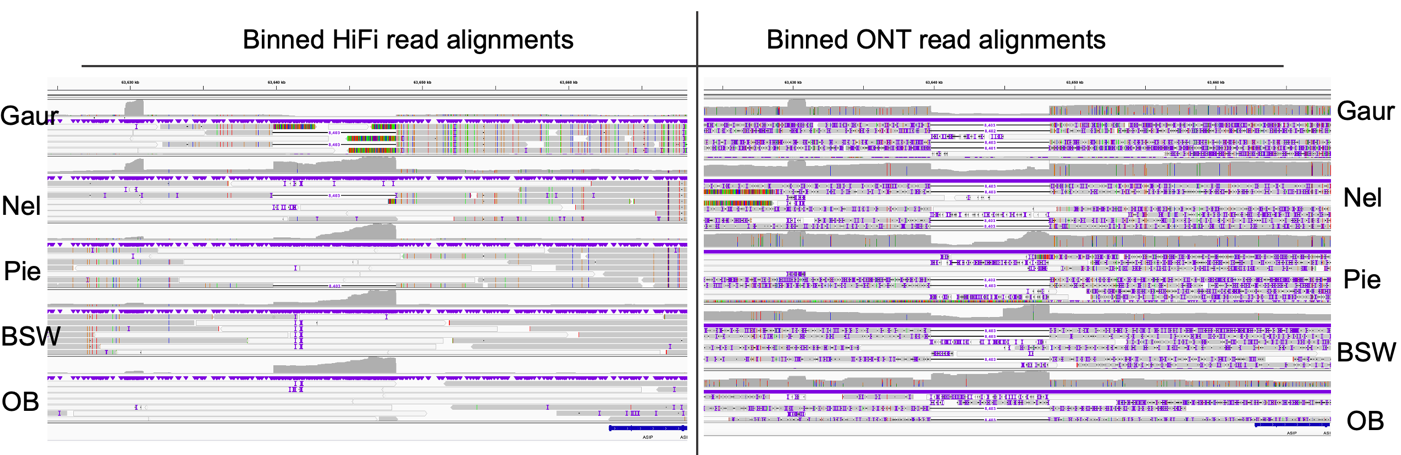

Figure SN_1_4: Binned HiFi (left) and ONT (right) read alignments against ARS-UCD1.2 indicate various structural rearrangements in the promoter of ASIP.

**Expansion of the coding sequence of *QRICH2* relative to ARS-UCD1.2**

A pangenome bubble on chromosome 19 overlapped with the fifth exon of *QRICH2* (ENSBTAG00000030173) encoding glutamine rich 2. Inspection of haplotype-resolved alignments revealed various insertions of 30 bp, 90 bp and 120 bp sequence at three distinct locations: 55,423,428 bp, 55,423,639 bp and 55,423,643 bp (Figure SN_1_5). At 55,423,428, an insertion of a nucleotide sequence «CCAGGCTGGTGTGGTCCAGCCCGGTGCAGC» is visible once in the Pied, BSW and OB assemblies, three times in Nellore, and four times in gaur. Additionally, the same sequence is inserted three times at 55,423,639 bp in the Nellore assembly. In the gaur sequence, a sequence of «CTGGTGCGGTGCAGCCCGGTGCAGGCCAGC» is inserted once at 55,423,642. This expansion of the coding sequence by multiples of 30 bp results in the addition of 10, 50, and 60 amino acids to the glutenine high molecular subunit of bovine QRICH2.

An amino acid motif «QxGxxQPxxx» occurs 16 times between amino acids 713 and 872 in the annotated coding sequence of bovine QRICH2 (ENSBTAT00000065208.2). This motif occurs 17, 21 and 22 times in the haplotype resolved taurine, Nellore, and gaur assemblies. A multi-species alignment (Sievers & Higgins, 2018) of different bovine QRICH2 versions with human orthologs confirms that this amino acid motif is also found in human QRICH2 (Figure SN_1_6).

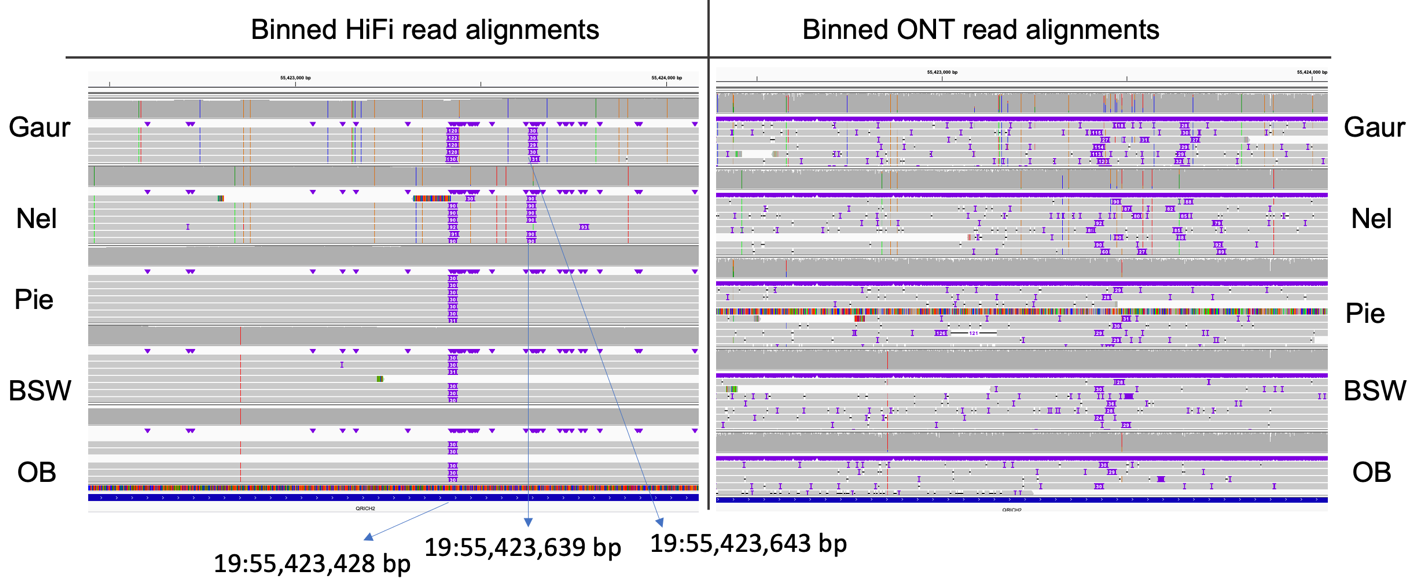

Figure SN_1_5: Binned HiFi (left) and ONT (right) read alignments against ARS-UCD1.2 indicate multiple instances of duplicated 30 bp sequence in the coding sequence of QRICH2.

Due to the highly repetitive character of the surrounding sequence, the insertion is difficult to discover from short read alignments, as it merely results in increased heterozygosity and allelic imbalance at heterozygous sites (Figure SN_1_7). The three insertion locations (55,423,428 bp, 55,423,639 bp and 55,423,643 bp) are readily identified by DeepVariant, but only as SNPs, with the first two having lower support than the reference base. Small or structural variant callers, like manta (Chen et al., 2016) and delly (Rausch et al., 2012), were also unable to identify any of the 30 bp insertions in any of the short read datasets. As such, this variation identified through de novo assemblies and pangenome integration appears completely opaque to short read analysis.

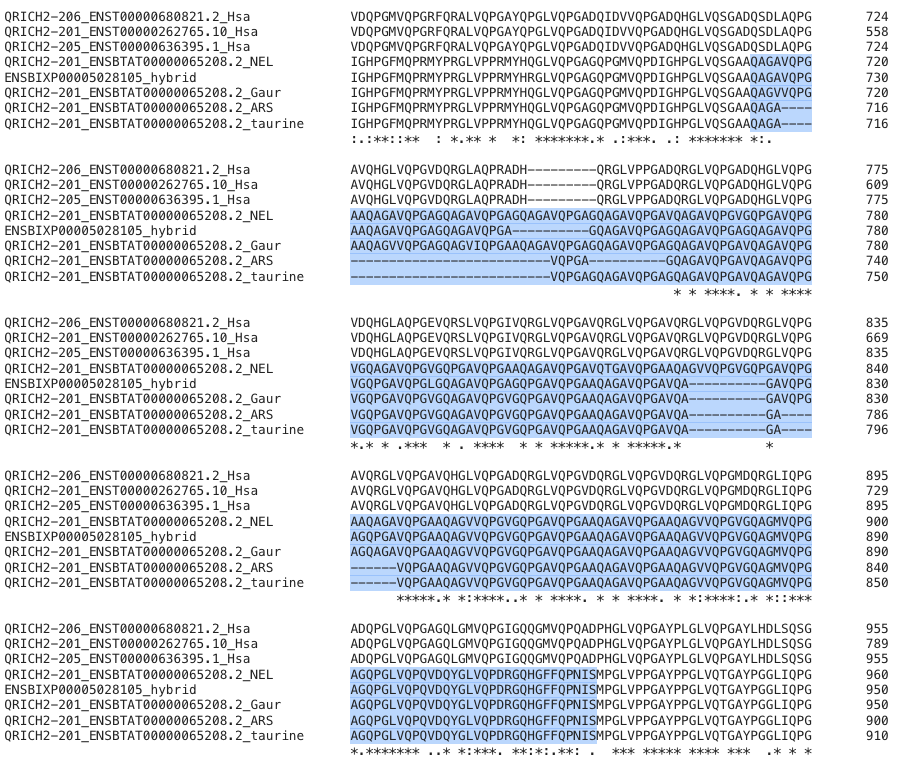

Figure SN_1_6: multi-species alignment of QRICH2. Alignment of the different versions of QRICH2 with its human orthologs. Blue background indicates amino acids 713 – 872 of bovine QRICH2 (ENSBTAT00000065208.2).

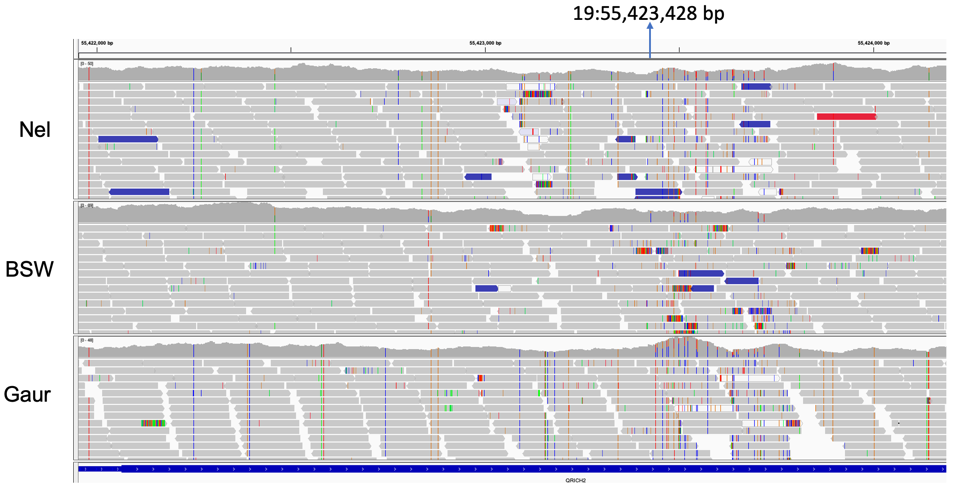

Figure SN_1_7: Short-read alignments against ARS-UCD1.2 at the fifth exon of QRICH2. Alignments of short reads from the Nellore sire and BSW dam of NxB, as well as from the gaur sire of GxP against ARS-UCD1.2 at the position where the long read alignments revealed the first insertion of 30 bp sequence.

Using RNA sequencing data from testis tissue of 76 mature taurine bulls, we confirm that *QRICH2* is transcribed in high abundance (31.52 ± 7.25 transcripts per million) in bovine testes. The fifth exon affected by the expansion of the coding sequence is also regularly expressed (Figure SN_1_8). However, a drop in sequencing coverage and many soft-clipped reads are visible at the region that harbors the repetitive motif likely indicating that increased ambiguity and incomplete reference sequence impairs read mapping (Figure SN_1_9). The raw RNA sequencing data used to investigate transcript abundance is available via (Kadri et al., 2021) from the European Nucleotide Archive at primary accession via PRJEB46995.

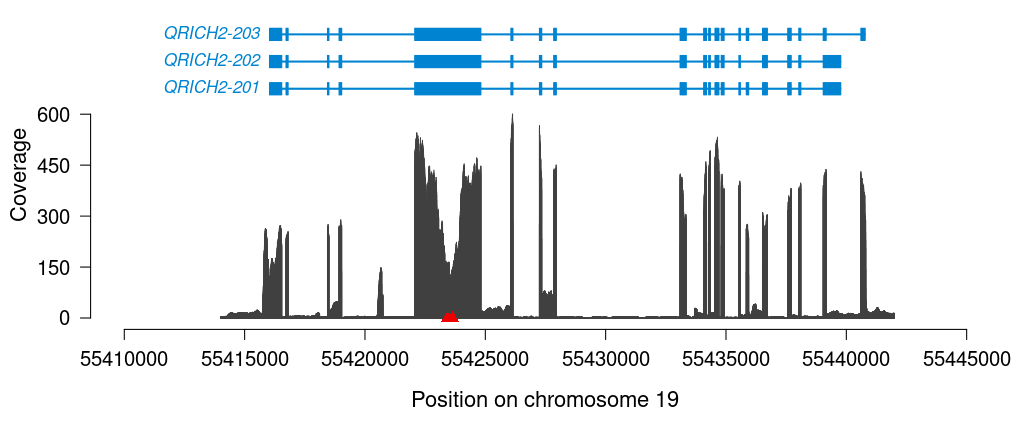

*Figure SN_1_8: Average coverage of RNA sequencing reads in testes of 76 mature bulls. The red diamonds represent the locations (55,423,428 bp, 55,423,639 bp, 55,423,643 bp) of the inserted sequences affecting the coding sequence of the fifth exon. Blue color represents the exon-intron structure of three* QRICH2 *isoforms.*

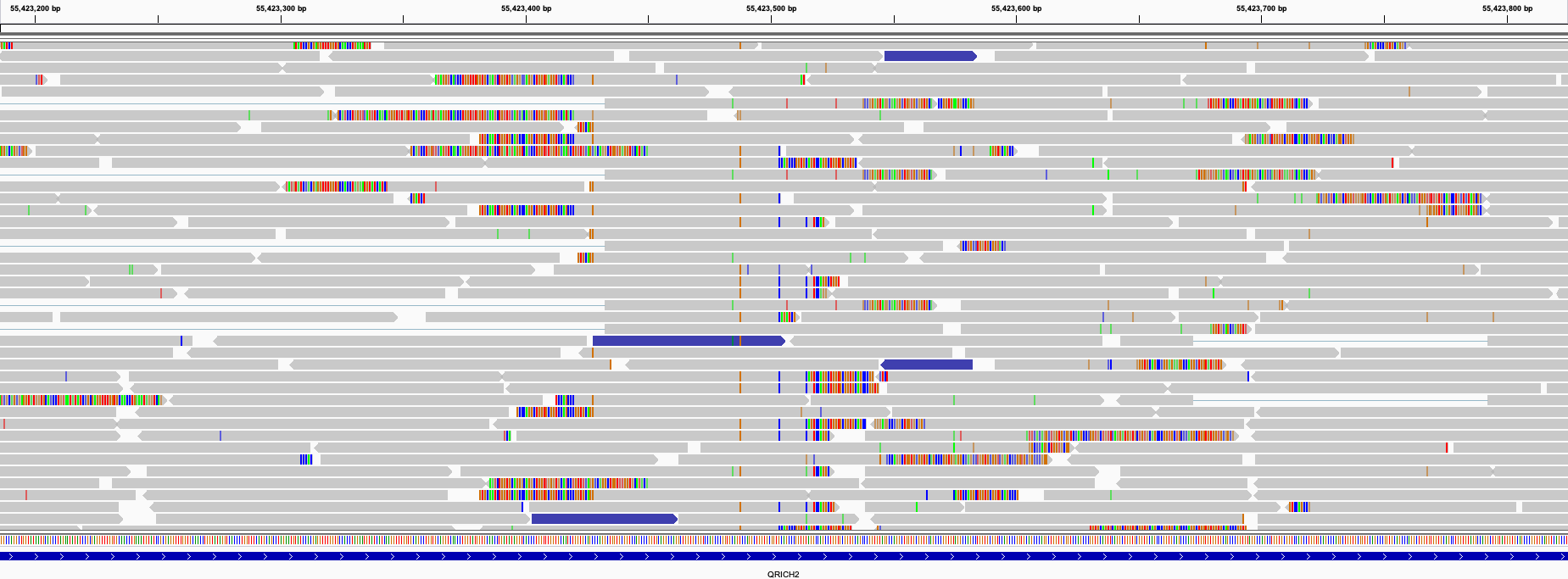

*Figure SN_1_9: Representative IGV screenshot of RNA sequencing data from testis tissue of a Brown Swiss bull aligned against ARS-UCD1.2 reference sequence at a region (Chr19:55,423,200 – 55,423,800) within the fifth exon of* QRICH2*. Many reads with soft clipped bases are indicative for the expansion of the coding sequence compared to ARS-UCD1.2. The raw RNA sequencing data underpinning this alignment are available at accession SAMEA9540538.*

Long read alignments of other publicly available long read bovid samples reveal additional expansions relative to ARS-UCD1.2 for yak, water buffalo, bison, and Highland cattle, but not in Simmental cattle (Figure SN_1_10). The presence of multiple insertions relative to ARS-UCD1.2 and other taurine assemblies in different bovid species suggests loss of *QRICH2* coding sequence in the taurine lineage. However, the error-prone CLR and ONT alignments are unable to cleanly resolve this challenging region. Alignments of long reads from Dominette, the cow on which ARS-UCD1.2 is based, indicate she may have carried the putative 30 bp deletion in the heterozygous state, or could have been an assembler/polishing collapse due to the high sequence similarity of these repeats. High-quality assemblies and long read alignments from additional taurine breeds and distantly related bovid samples are required to clearly disentangle the evolutionary relationship surrounding the *QRICH2* SVs.

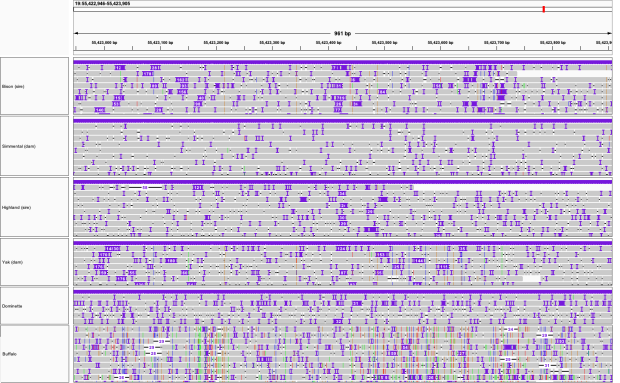

Figure SN_1_10. SVs in QRICH2 across other bovids. Alignments of trio binned ONT reads from a bison x Simmental cross (Heaton et al., 2021; Oppenheimer et al., 2021), trio binned CLR reads from a Highland x yak (Rice et al., 2020) cross, and CLR reads from a water buffalo (Low et al., 2019) reveal similar patterns of insertions relative to ARS-UCD1.2. Alignments of CLR reads from Dominette (Rosen et al., 2020) also indicate a potential insertion at the expected coordinate.

**Deletion of *TAS2R46* and *ENSBTAG00000001761***

A pangenome bubble on chromosome 5 overlapped with *TAS2R46* (ENSBTAG00000030471) and *ENSBTAG00000001761* encoding taste 2 receptor member 46 and olfactory receptor family 6 subfamily AA member 1, respectively. Inspection of long read alignments confirmed deletion of 17,017 bp in the gaur assembly between 98,587,384 and 98,604,401 bp encompassing the entire coding sequences of *TAS2R46* and ENSBTAG00000001761 (Figure SN_1_11). This deletion was not detected in the other haplotype-resolved assemblies.

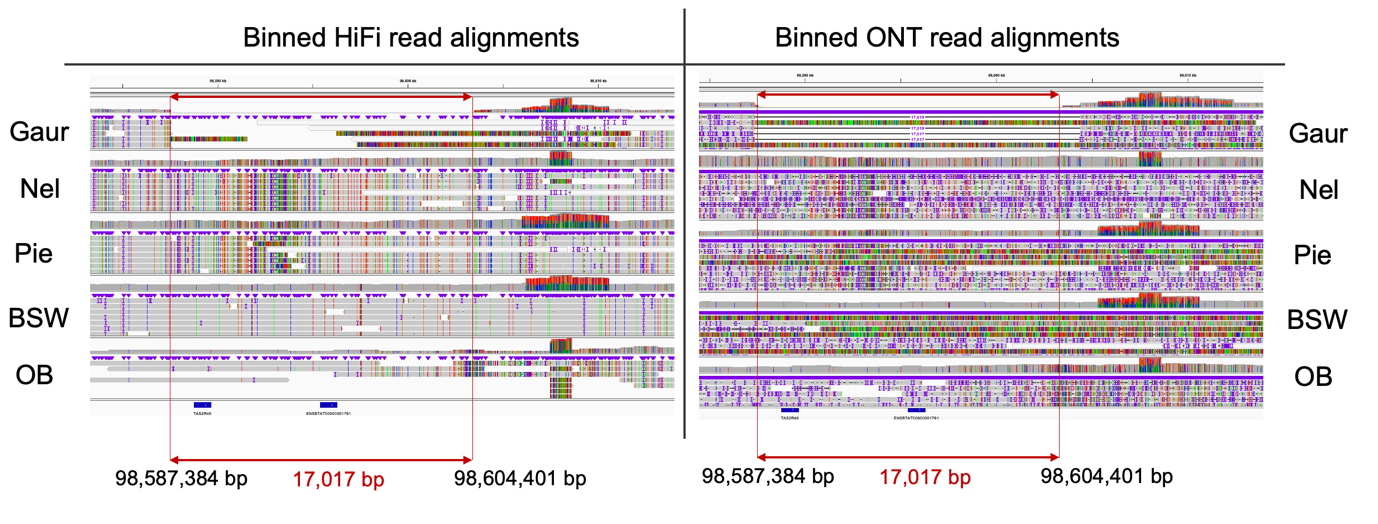

Figure SN_1_11: Binned HiFi (left) and ONT (right) read alignments against ARS-UCD1.2 indicate deletion of 17 kb in the gaur assembly. Red color highlights the coordinates of the deletion in gaur encompassing two genes (blue tracks at the bottom).

Coverage of short read alignments from the gaur sire verifies the presence of the deletion (Figure SN_1_12). The gaur sire carries the deletion in the heterozygous state suggesting within-species variability and the transmission of the haplotype carrying the deletion to the F1.

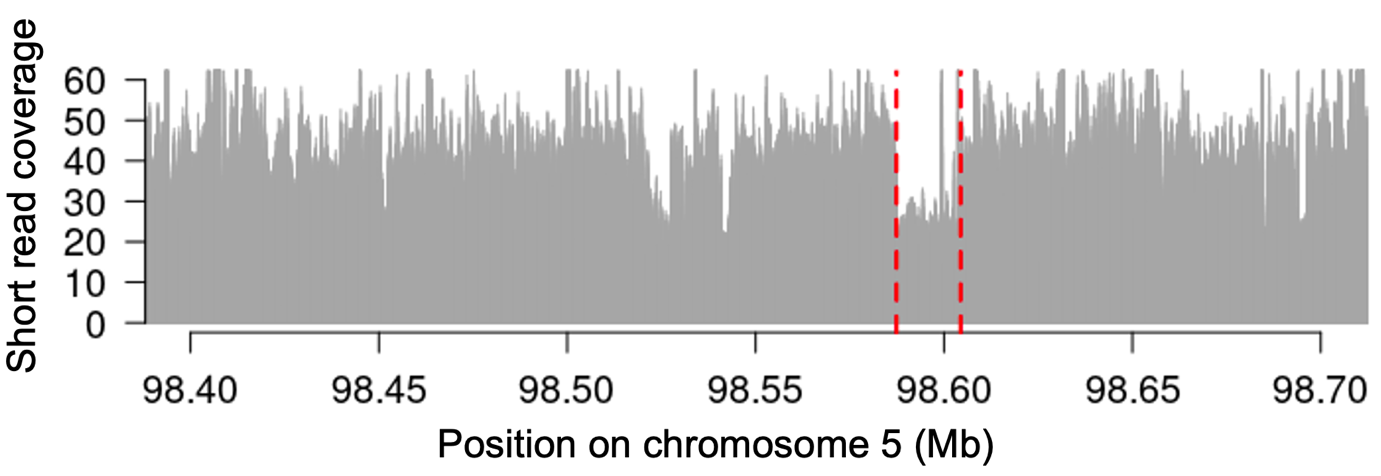

Figure SN_1_12: Short read alignments from the gaur sire against ARS-UCD1.2 confirm the deletion of a 17 kb segment on chromosome 5. Red dotted lines indicate the location of deletion. Short read coverage at the deletion is approximately half the average coverage, indicating that the gaur is heterozygous carrier of the deletion.

**Duplication of *HSPA1A***

A pangenome bubble at chromosome 23 overlapped *HSPA1A* (ENSBTAG00000025441) encoding heat shock protein family A (Hsp70) member 1A. We observed elevated coverage at a 2,294 bp segment between 27,520,653 bp and 27,522,947 bp and multiple soft-clipped reads extending the seemingly duplicated region in all haplotype-resolved alignments (Figure SN_1_13). The region of elevated coverage encompasses the 5’UTR and the entire exon of *HSPA1A*. Manual inspection and assembly of the soft clipped bases enabled reconstructing an insertion of approximately 11 kb insertion (Figure SN_1_14). The 11 Kb insertion can also be identified through calling structural variants with the long reads, as confirmed by pbsv (https://github.com/pacificbiosciences/pbsv). Elevated coverage at the 2,294 bp segment encompassing *HSPA1A* is likely due to reads from *HSPA1B* that cannot align properly, as *HSPA1B* is missing in the current *Bos taurus* reference genome (Suqueli García et al., 2017). The pangenome region and respective repeat analysis is shown in Figure SN_1_15.

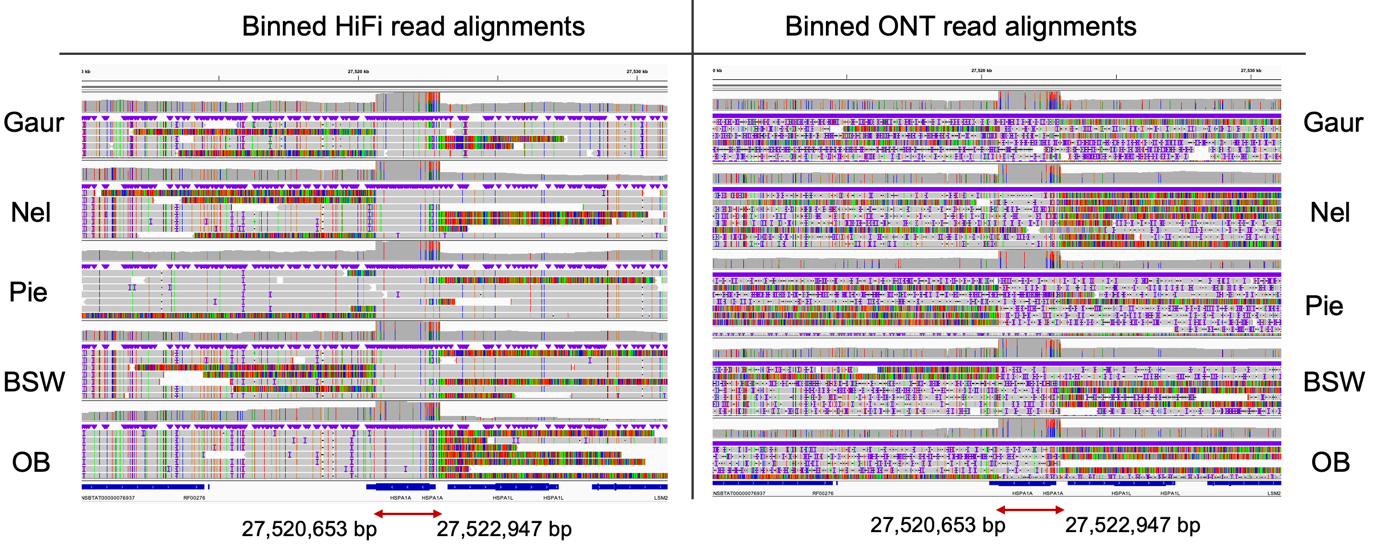

Figure SN_1_13: Binned HiFi (left) and ONT (right) read alignments against ARS-UCD1.2 as well as the corresponding coverage tracks indicate duplication of a 2.3 kb segment encompassing HSPA1A in all assemblies.

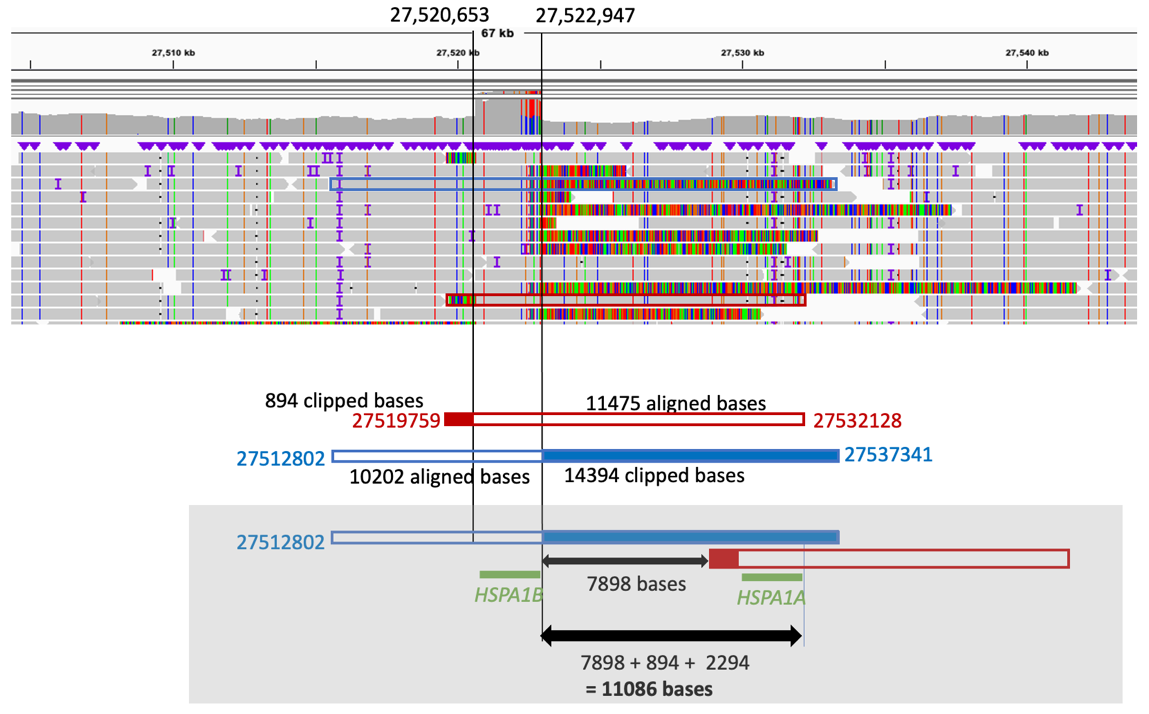

Figure SN_1_14: Recovery of an approximately 11 kb insertion in the Nellore haplotype from manual assembly of reads with soft-clipped bases.

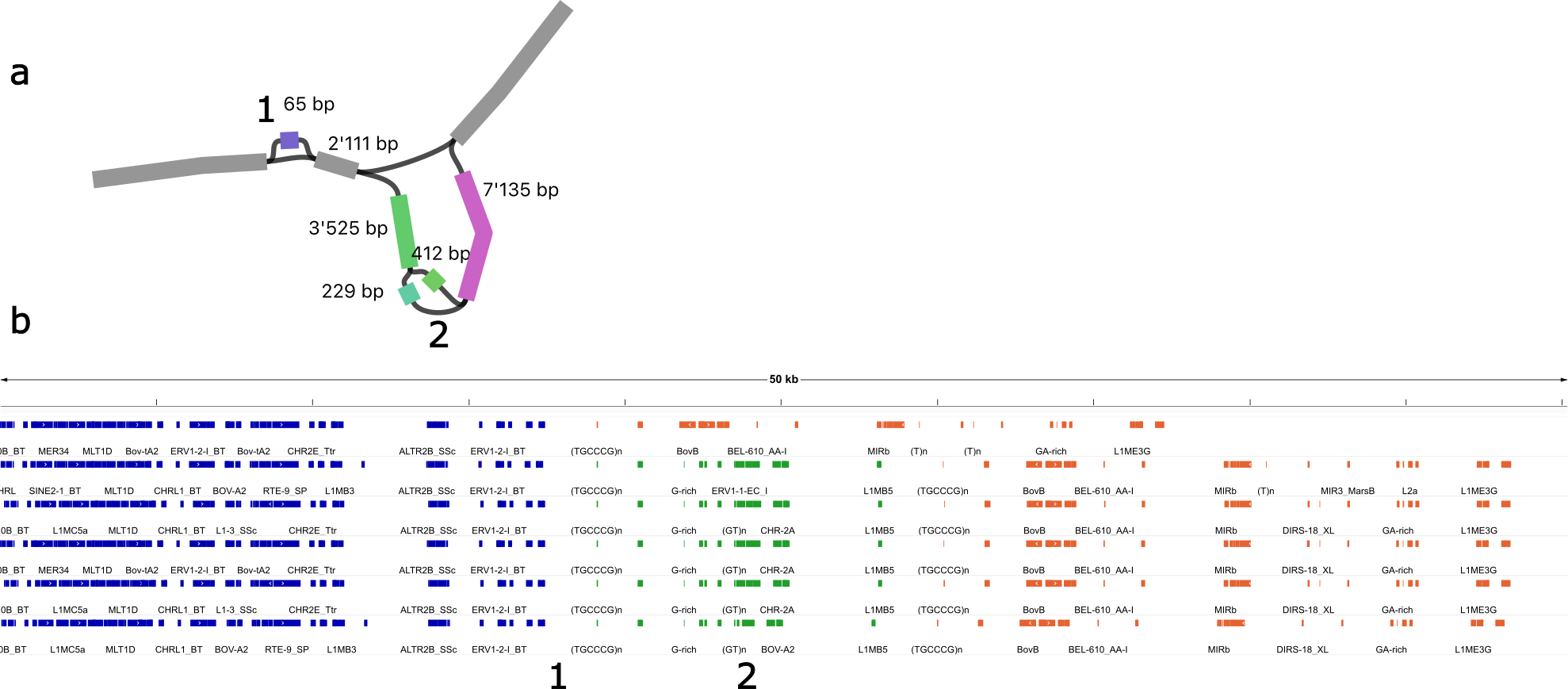

Figure SN_1_15: The hifiasm pangenome region around HSPA1A (located within the orange repeats) predicts an 11 Kb insertion, which contains HSPA1B (located within the green repeats). The Original Braunvieh assembly contains a 65 bp insertion (marked as “1”), while the gaur assembly contains a 200 bp deletion (marked as “2”). The hifiasm gaur haplotype path through the insertion is approximately 10,889 bp, while the other haplotype path is 11,072 bp, in strong agreement with the soft-clipped read estimation from Figure SN_1_14.

**Insertion and deletion of 84 bp in the coding sequence of *PRDM9***

The pangenome graph contained a bubble on chromosome 1 with an 84 bp deletion that overlapped the eleventh exon of *PRDM9* (ENSBTAG00000004538) encoding PR domain containing 9. Read alignments confirm the deletion of 84 bp («GCCCACACTCCCCGC	AAACATAGGGCTTCTCCCCTGTGTGTGTCCTCTGGTGTGTGATGAGAGTGGACTTCTGATTGAAGCTTT») between 157,543,126 bp and 157,546,209 bp in the gaur and Nellore assemblies (Figure SN_1_16). In the paternal Original Braunvieh haplotype, an insertion of 84 bp («CCCACACTCCCCGCAAACATAGGGCTTCTCCCCTGTGTGTGTCCTCTGGTGTGTGATGAGACGGCCCTTCTGACTGAAGCTTTG») occurred at 157,546,378 bp.

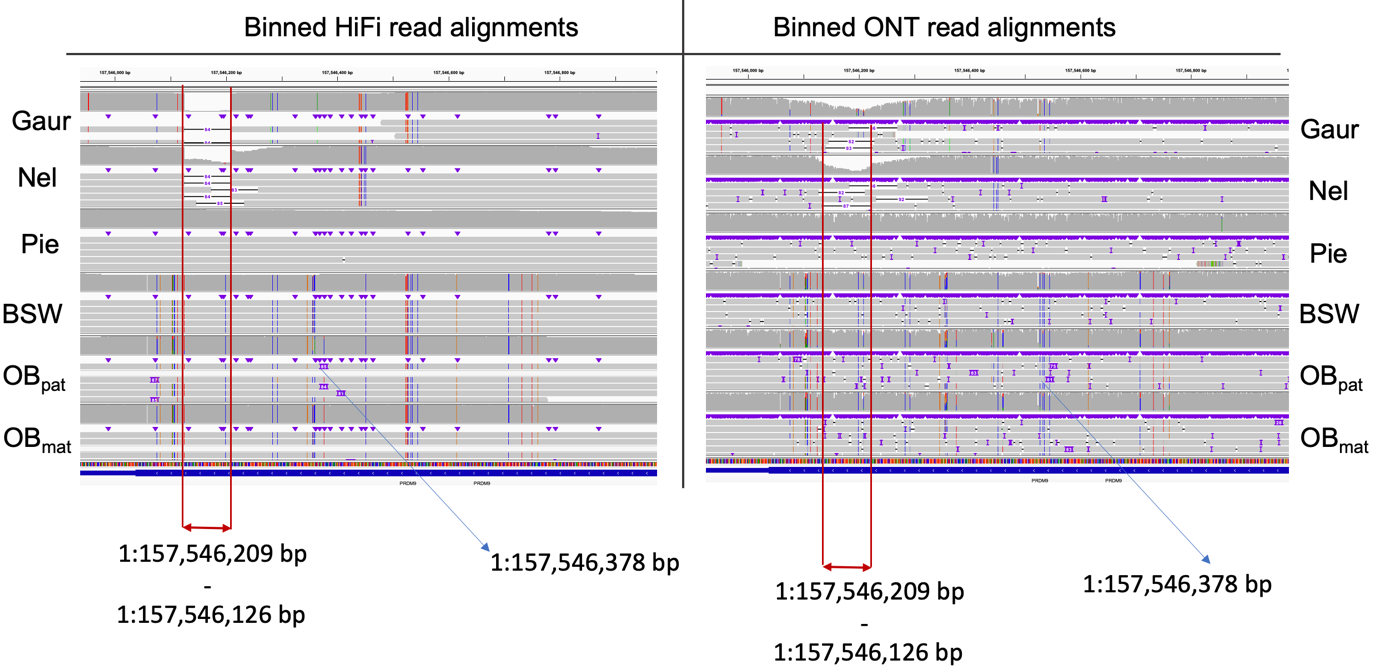

Figure SN_1_16: Binned HiFi (left) and ONT (right) read alignments against ARS-UCD1.2 as well as the corresponding coverage tracks indicate deletion and insertions of 84 bp in the eleventh exon of PRDM9.

Due to the repetitive nature of the sequence encoding the zinc fingers, this variation is difficult to resolve using short read alignments (Figure SN_1_17).

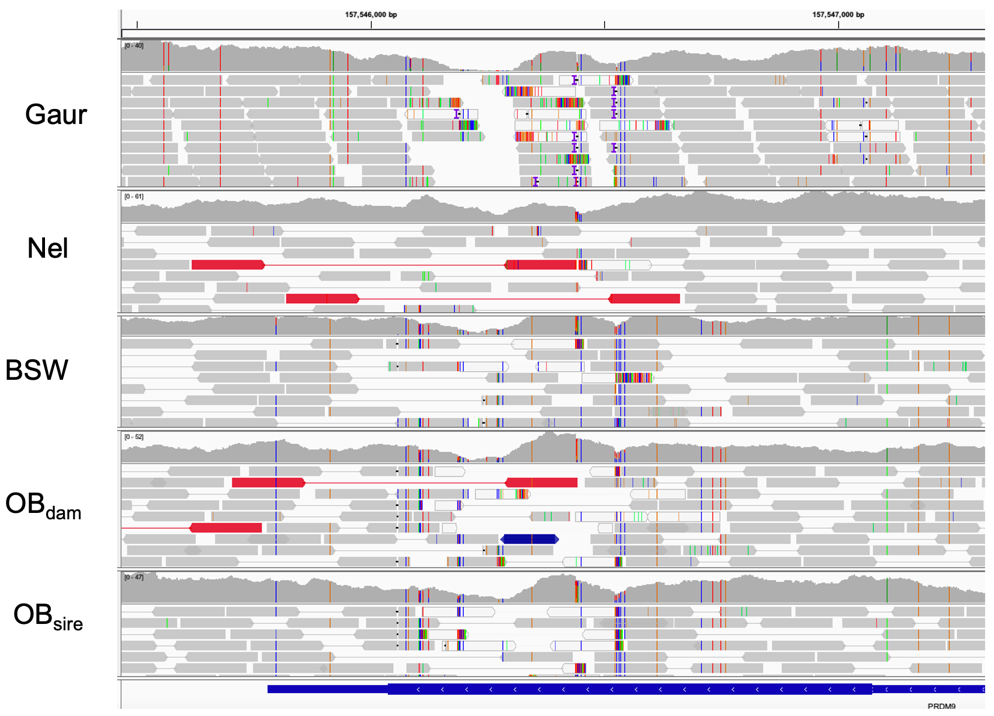

Figure SN_1_17: Alignments of parental short reads against ARS-UCD1.2 at the eleventh exon of PRDM9. Red colored symbols and low coverage indicate the deletion in the gaur and Nellore sires.
